# Uptake of Fe-fraxetin complexes, an IRT1 independent strategy for iron acquisition in *Arabidopsis thaliana*

**DOI:** 10.1101/2021.08.03.454955

**Authors:** Kevin Robe, Max Stassen, Joseph Chamieh, Philippe Gonzalez, Sonia Hem, Véronique Santoni, Christian Dubos, Esther Izquierdo

## Abstract

- Iron (Fe) is a micronutrient essential for plant growth and development. Iron uptake in alkaline soil is a challenge for most plants. In this study, we investigated the role of the catechol coumarins fraxetin and esculetin in plant Fe acquisition and their Fe chelating properties.
- Mass spectrometry and capillary electrophoresis were used to characterize Fe-coumarin complexes. To understand the role of these complexes, genetic, molecular and biochemical approaches were deployed.
- We demonstrated that catechol coumarins are taken up by *Arabidopsis thaliana* root via an ATP dependent mechanism and that plants defective in IRT1 activity (the main high affinity Fe importer) or bHLH121 (a key regulator of Fe deficiency responses) can be complemented by exogenous supply of fraxetin and to a lesser extent of esculetin. We also showed that Fe and fraxetin can form stable complexes at neutral to alkaline pH that can be taken up by the plant.
- Overall, these results indicate that at high pH, fraxetin can improve Fe nutrition by directly transporting Fe(III) into the root, circumventing the FRO2/IRT1 system, in a similar way as phytosiderophores do in grasses. This strategy may explain how non-grass species can thrive in alkaline soils.

## INTRODUCTION

Iron (Fe) is a major micronutrient required for plant growth and development whose availability affects crop productivity and the quality of their derived products (Briat *et al*., 2015). Although Fe is one of the most abundant elements in soil, plants often suffer from Fe deficiency (Chen & Barak, 1982), which most often occurs in alkaline soils, where Fe solubility is low. In aerated soils, Fe solubility decreases by 1000 times for each pH unit increase between 4 and 9 (Lindsay, 1979). The poor Fe bioavailability in soil has led plants to evolve strategies for Fe mobilisation and uptake, the so called Strategy I and Strategy II (Gao & Dubos, 2021).

Grass species (Strategy II plants) release Fe mobilising compounds, called phytosiderophores, that form stable complexes with both Fe(III) (ferric iron) and Fe(II) (ferrous iron). Phytosiderophore-Fe complexes are then taken up into the plant by the epidermis-localised transporter YS1 (YELLOW STRIPE 1) family (Curie *et al*., 2001; Weber *et al*., 2006; Xuan *et al*., 2006). When it is expressed in the Fe uptake-defective yeast mutant *fet3fet4*, YS1 can restore the growth of yeast grown in a phytosiderophore-containing medium, even in the presence of the Fe(II) chelator BPDS (bathophenanthroline disulfonic acid) (Meda *et al*., 2007). It is noteworthy that other micronutrients than Fe, such as copper (Cu), zinc (Zn), nickel (Ni) and cobalt (Co), also form complexes with phytosiderophores and are taken up as such by the plant (Xuan *et al*., 2006).

In non-grass species (Strategy I), efficient Fe uptake from soil is ensured by a reduction-based mechanism (Kobayashi *et al*., 2012; Brumbarova *et al*., 2015). This process involves the acidification of the rhizosphere through the release of protons (H^+^), the reduction of Fe(III) to Fe(II) by a membrane localised reductase and the subsequent transport of Fe(II) across the rhizodermis cell plasma membrane by high affinity Fe transporters. In *Arabidopsis thaliana*, these three steps rely on the activity of AHA2 (ATPase2), FRO2 (FERRIC REDUCTION OXIDASE2) and IRT1 (IRON-REGULATED TRANSPORTER1), respectively (Robinson *et al*., 1999; Vert *et al*., 2002).

At alkaline pH, FRO2 activity is negligible (Terés *et al*., 2019) and as a consequence, Fe(III) cannot be effectively reduced, leading to low Fe uptake. To cope with the inactivity of FRO2 under high pH, Fe-mobilising coumarins (i.e. catechol coumarins such as esculetin, fraxetin and sideretin in *Arabidopsis thaliana*) are secreted in the rhizosphere by the ABCG transporter PDR9 (PLEIOTROPIC DRUG RESISTANCE 9) (Fourcroy *et al*., 2014). Coumarins are secondary metabolites that derive from the phenylpropanoid pathway (Rodríguez-Celma *et al*., 2013; Fourcroy *et al*., 2014; Schmidt *et al*., 2014). In *Arabidopsis thaliana*, Feruloyl-CoA 6-Hydroxylase1 (F6H1), a 2-oxoglutarate-dependent dioxygenase enzyme, is responsible for the ortho-hydroxylation of feruloyl-CoA into 6-hydroxyferuloyl-CoA, the common precursor of coumarins (Rodríguez-Celma *et al*., 2013; Schmid *et al*., 2014). Because scopoletin lacks catechol groups, it cannot bind to Fe and thus is not directly involved in iron acquisition (Rajniak *et al*., 2018). The two main catechol coumarins fraxetin and sideretin are biosynthesised by S8H (SCOPOLETIN 8-HYDROXYLASE), a 2-oxoglutarate-dependent dioxygenase (Siwinska *et al*., 2018; Tsai *et al*., 2018) and CYP82C4 P450-dependent monooxygenase, respectively. Both genes have been shown to be upregulated by iron deficiency (Murgia *et al*., 2011; Rajniak *et al*., 2018). While alkaline pH favours the biosynthesis of fraxetin, the main catechol coumarin secreted at acidic pH is sideretin. The importance of catechol coumarin secretion for Fe nutrition has been demonstrated in several studies (Schmid *et al*., 2014; Rajniak *et al*., 2018; Tsai *et al*., 2018). It has been shown that catechol coumarins can reduce Fe(III) to Fe(II) and form stable complexes with Fe(III); however, the relative importance of each phenomenon for the plant and the precise function of these metabolites in iron nutrition is still not fully understood (Robe *et al*., 2021b). One might formulate three hypotheses on the role of catechol coumarins in plant Fe nutrition: (i) catechol coumarins form stable complexes with Fe(III) that are translocated into the roots through the activity of a yet unknown transporter localised at the root epidermis cells or (ii) the Fe(III) solubilised by coumarins is used as substrate for the FRO2 reductase, or (iii) the Fe(III) is reduced by the catechol coumarins and directly taken up by IRT1 (Rodríguez-Celma & Schmidt, 2013). This three hypotheses are not mutually exclusive and may co-exist. The uptake of catechol coumarins by Strategy I plants grown in Fe deficient conditions has already been reported (Robe *et al*., 2021a); however, the biological role of this phenomenon was still to be decrypted. These recent results led us to consider coumarins as putative siderophores. In the present work we show that, at circumneutral pH, the catechol coumarin fraxetin forms stable complexes with Fe(III) that are taken up by plant roots and in turn significantly improve Fe nutrition of mutants lacking the high affinity Fe(II) transporter IRTI and also in the presence of the strong Fe(II) chelator ferrozine. The ATP-dependent uptake of Fe-coumarin complexes by the plant root, in a similar way to phytosiderophores uptake by grasses, is a yet undescribed mechanism for Fe uptake in Strategy I plants. Taken together, these data further confirm that the boundaries between Strategy I and Strategy II plants are much narrower than what is classically described.

## MATERIALS AND METHODS

### Plant Material

*Arabidopsis thaliana* ecotype Columbia (Col-0) was used as wild type (WT). The *Arabidopsis thaliana* mutant lines used in this study are: *f6’h1-1(*Kai *et al*., 2008), *irt1-2* (Varotto *et al*., 2002), *fro2* (*frd1-1*) (Robinson *et al*., 1999) and *bhlh121-2* (Gao *et al*., 2020a; Gao *et al*., 2020b). The Fe-inefficient tomato (*Solanum lycopersicum*) mutant used in this study was T238*fer (*Brown et *al*., *1971)*

### Growth conditions

For *in vitro* culture, seeds were surface sterilised with a solution containing 12.5/37.5/50 (v/v/v) of bleach/water/ethanol for 5 minutes with strong agitation. Seeds were rinsed 3 times with 96% ethanol before drying. Seeds were then germinated on control Hoagland medium (50 µM Fe-EDTA at pH 5.5) solidified with 0.7% agar and grown under long day condition (16h light/8h dark).

For all the experiments control Hoagland medium contained 50 µM Fe-EDTA and poorly available Fe Hoagland medium contained 25 µM FeCl_3_, supplemented with 0.5 g l^-1^ HEPES buffer and adjusted to 7 with KOH.

For coumarin uptake, *f6’h1* mutant plants were grown in hydroponic condition for 4 weeks in control Hoagland solution. Coumarins were added for 3 days at 50 µM in a new Hoagland solution containing poorly available Fe (25 µM FeCl_3_ at pH 7). For orthovanadate and glibenclamide treatments, following 1 week of growth on control Hoagland, seedlings were transferred for another 3 days in Hoagland containing poorly available Fe and 100 µM sodium orthovanadate or 150 µM glibenclamide (both from Sigma-Aldrich). For agar plates, pH was adjusted after autoclave using a pH indicator paper (± 0.25 pH units). For coumarin uptake kinetics, one-week-old seedlings were transferred to Hoagland agar plates containing poorly available Fe and 50 µM fraxetin.

For *in vitro* mutant complementation, plants were directly germinated on poorly available Fe Hoagland plates (25 µM FeCl_3_ at pH 7) containing the synthetic coumarins scopoletin, fraxetin or esculetin (all three from Sigma-Aldrich). Scopoletin was added at a concentration of 25 µM, esculetin at 50 µM and fraxetin at 100 µM. 100 µM concentration for scopoletin or esculetin was not used since detrimental effects on seedlings were observed.

Tomato seeds were germinated for 1 week on wet tissue paper under long day condition. When cotyledons were fully developed, seedlings were transferred to a Hoagland solution containing 25 µM FeCl_3_ at pH 7.5, supplemented or not with 100 µM fraxetin. Plants were then grown under short day condition (8 h light/ 16h dark).

### Coumarin imaging

Seedlings were grown for one week on control Hoagland agar plates (50 µM Fe-EDTA pH 5,5) prior to their transfer for 3 days on low Fe availability medium (25 µM FeCl_3_, pH 7) supplemented with 25 µM of scopoletin, fraxetin or esculetin. The root cell walls were stained with 10 µg ml^-1^ propidium iodide (PI) during 10 min. Coumarins were imaged with an LSM 780 multiphoton (i.e. two photon) microscope (Zeiss, Oberkochen, Germany) equipped with a Chameleon Ultra II laser (Coherent, Santa Clara, CA, USA) and a W Plan Apochromat 20x 1.0 objective (Talamond et al., 2015; Koyyappurath et al., 2015). Spectral imaging was performed using laser excitation at 720 nm and the 32-channel GaAsP spectral detector as described elsewhere (Robe *et al*., 2021a). The Linear Unmixing function in Zeiss Zen microscope software (version 2.10) was used to separate the signals of the auto-fluorescent compounds scopolin, fraxin, esculin and PI.

### Chlorophyll Content

Chlorophyll from 2 to 5 mg of leaf tissue (fresh weight) were extracted in 1 ml of 100% acetone in the dark under agitation. The absorbance (A) at 661.8 and 644.8 nm was then measured. Total chlorophyll content was assessed using the following equation: Chl a + Chl b = 7.05 x A_661.6_ + 18.09 x A_644.8_ and was expressed as micrograms per gram fresh weight.

### HPLC analysis of roots and leaves extracts

Coumarin extraction from roots, samples preparation for HPLC and coumarin quantification were performed as described previously (Robe *et al*., 2021a). HPLC-UV and HPLC-fluorescence analysis were performed using a 1220 Infinity II LC system (Agilent Technologies) coupled to a Prostar 363 fluorescence detector (Varian). Separation was done on an analytical HPLC column (Kinetex 2.6 µm XB-C18 100 Å, 150 × 4.6 mm; Phenomenex), with a gradient mobile phase made with 0.1% (v/v) formic acid in water (A) and 0.1% (v/v) formic acid in acetonitrile (B) and a flow rate of 0.5 ml min^-1^. The gradient program started at 8% B for 1 min and increased linearly to 33% B in 11.5 min and then to 50% B in 0.5 min. This proportion was maintained for 3 min and returned linearly to initial conditions in 0.5 min. The column was allowed to stabilise for 5 min at the initial conditions. Absorbance was monitored at λ = 338 nm. Fluorescence was monitored at λexc 365 and λem 460 nm

### LC-MS/MS analysis of soil grown roots

Wild type and *f6’h1* plants were co-cultivated for three weeks in soil supplemented with 0 to 0.6% (w/w) CaO. Plants were grown at a distance sufficient to prevent root mixture between genotypes. Soils with growing Arabidopsis plants were roughly washed with tap water. Roots of plants were then gently rinsed with distilled water until removing all soil particles. Then coumarins were extracted from roots as previously described for HPLC analysis (Robe *et al*., 2021a).

LC-MS/MS analysis were performed using an Ultimate 3000 nano HPLC system (Thermo Fisher Scientific Inc.) coupled to a quadrupole time-of-flight (Q-TOF) mass spectrometer (Maxis Impact; Bruker Daltonics GmbH) as described previously (Robe *et al*., 2021a). The LC conditions were those used for HPLC analysis with the exception of the flow rate, which was reduced to 200 µl min^−1^. Glycosylated coumarins usually lose their sugar moiety in the ESI source. Therefore, after the verification of their identity by retention time, MS and MS/MS profiles, fraxin, esculin, scopolin and sideretin glycosides were quantified by integrating the MS Extracted Ion Chromatograms (EIC) peaks of their aglycone counterparts. As no standards are commercially available for sideretin glycosides, calibration curves could not be performed and thus, relative quantification was done for all the coumarins in this experiment. Results are expressed as relative intensity peak areas (cts min^-1^).

### Direct infusion mass spectrometry

To study the *in vitro* interaction of Fe and other transition metals with coumarins, 50 µM methanolic solutions of each ligand (fraxetin, scopoletin and esculetin) and 50 µM aqueous solution of each metal salt (FeCl_3_, MnCl_2_, ZnCl_2_, NiCl_2_, AlCl_3_ and CuCl_2_) were freshly prepared. Mixtures of metal:ligand ratios of 1:2 were made and diluted to 1/1000 in a solution of 50% acetonitrile/50% 25 mM ammonium bicarbonate (adjusted to pH 8, 7, 6, 5 or 3 by addition of formic acid). The obtained solutions were injected into the mass spectrometer via a syringe pump at a rate of 200 µl h^-1^. Source settings were the same as used for LC-MS analysis except for mass range: 50 to 1000 m/z and ESI that was operated in negative mode.

### Capillary electrophoresis and Taylor dispersion analysis

Capillary electrophoresis experiments were performed on an Agilent 7100 capillary electrophoresis instrument with a built-in diode array UV detector (Waldbronn), detection was performed at 214 nm. Fused silica capillaries of 50/375 µm I.D./O.D. were from Polymicro Technologies. Capillary dimensions were 33.5 cm long (25 cm to detection window). New capillaries were conditioned by performing the following washes at 1 bar: 1 M NaOH for 30 min and water for 15 min. Before each sample, the capillary was flushed with the buffer for 2 min. Samples were injected hydrodynamically on the inlet side of the capillary by applying a 50 mbar pressure for 3 s. Separations were achieved by applying a -15 kV voltage. Injections were repeated 3 times, between each repetition a 30 s flush with the buffer was done. Samples were prepared at a 1 µM concentration of fraxetin in a 50 mM ammonium bicarbonate (NH_4_HCO_3_) buffer at variable pH (5 to 8) adjusted with formic acid or in a 50 mM formic acid solution pH 2.5.

The hydrodynamic radius of fraxetin was determined using Taylor dispersion Analysis (Chamieh & Cottet, 2014). Briefly, using the same capillary, instrument and buffers as for the capillary electrophoresis experiments, the samples were injected hydrodynamically by applying a 50 mbar pressure for 5 s and were mobilised with the buffer by applying a 100 mbar pressure. The elution profiles were treated as described elsewhere (Chamieh & Cottet, 2014) to calculate the diffusion coefficient and the hydrodynamic radius (*R*_*h*_). In the case of Fe(III)-fraxetin complexes, the *R*_*h*_ was estimated from the diffusion coefficient obtained using HydroPro software (Ortega *et al*., 2011) on structures drawn with Marvin Sketch (ChemAxon).

The determination of the effective charge of fraxetin and Fe-fraxetin complexes was achieved by using the method based on the O’Brien-White-Ohshima model (OWO) (Ohshima, 2001; Ibrahim *et al*., 2013). Protocols and equations for the determination of the effective charge of fraxetin and Fe(III)-fraxetin complex and the pKa of Fraxetin are detailed in Supporting Information Methods S1.

### Metals measurements

About 2.5 mg of dry root or 7.5 mg of dry shoot tissue per sample were mixed with 750 ml of nitric acid (65% [v/v]) and 250 ml of hydrogen peroxide (30% [v/v]). After one night at room temperature, samples were mineralised at 85°C during 24 hours. Once mineralised, 4 ml of milliQ water was added to each sample. Metal contents present in the samples were then measured by microwave plasma atomic emission spectroscopy (MP-AES, Agilent Technologies).

### Western blot analysis

Western blots were performed as described previously (Robe *et al*., 2021a). Imunodetection of ferritins was performed using a rabbit polyclonal antiserum raised against purified FER1 (Dellagi *et al*., 2005). Dilutions of antibodies applied were: rabbit anti-ferritin 1:10000 (primary) and anti-rabbit HRP conjugated (Promega) 1:10000 (secondary).

### Accession Numbers

Sequence data from this article can be found in the GenBank/EMBL libraries under the following accession numbers: *F6’H1* (AT3G13610), *IRT1* (AT4G19690), *FRO2* (AT1G01580), *bHLH121* (AT3G19860).

## RESULTS

### Coumarin uptake in the presence of iron is an active mechanism

The uptake and glycosylation of coumarins by Arabidopsis roots under Fe deficient conditions was previously shown (Robe *et al*., 2021a). When *f6’h1* mutants (impaired in Fe mobilising coumarin biosynthesis; Fig. **1a**) are grown in close vicinity to wild type (WT) plants in agar plates without any Fe, the coumarins secreted by WT into the medium (scopoletin, fraxetin and esculetin) are taken up by *f6’h1* roots. Uptaken coumarins are then glycosylated and stored in the root vacuoles in the form of their glycosylated counterparts (scopolin, fraxin and esculin) (Robe *et al*., 2021a). However, this mechanism has never been assessed for plants grown in natural soil, where large amounts of insoluble iron hydroxides are expected to accumulate. This was assayed using *f6’h1* mutant plants grown in close vicinity to WT plants in potting soil supplemented with different calcium oxide (CaO) concentrations (i.e. 0%, 0.2%, 0.4% and 0.6%). CaO was used to increase the pH of the soil and thus to stimulate the production of coumarins such as fraxetin and esculetin (Rajniak *et al*., 2018). Since, in roots, coumarins are mostly present in their glycosylated form, only coumarin glucosides where quantified in this tissue. As no commercial standards are available for sideretin glucosides, we used mass spectrometry (MS) to unequivocally identify the coumarins that are produced and taken up (Fig. **1b**). As expected, in WT plants, scopoletin and fraxin accumulation was increased in a CaO concentration dependent manner whereas the accumulation of sideretin glucoside followed the opposite trend. In contrast, none of the studied coumarins were identified in the roots of the *f6’h1* mutant grown alone for all the CaO concentration tested. However, MS analysis of *f6’h1* roots co-cultivated with WT confirmed that fraxetin and the non-catechol coumarin scopoletin can be taken up by Arabidopsis root when grown in soil. Esculin content in this experiment was very low and it could not be quantified by MS. Sideretin glucosides were detected in *f6’h1* roots co-cultivated with WT and not in *f6’h1* roots grown alone, showing that sideretin can also be taken up by plant roots (Fig. **1b**).

According to their octanol-water partition coefficient (Log K_ow_) values (∼1.5) (Mercer *et al*., 2013), the coumarins studied here are predicted to diffuse through plant roots in a passive manner (Limmer & Burken, 2014; Selmar *et al*., 2015). However, the inability of rice and maize to take up fraxetin (Robe *et al*., 2021a) suggests that the uptake of catechol coumarins is not passive and is mediated by a transporter only present in Strategy I plants. We investigated the uptake of coumarins using sodium orthovanadate (Na_3_VO_4_), an inhibitor of plasma membrane ATPases that inhibits ABC transporter activities (Kang *et al*., 2011). For this purpose, *f6’h1* mutant was grown in hydroponic condition and supplemented with two non-catechol coumarins (scopoletin and umbelliferone) and two catechol coumarins (fraxetin and esculetin) in the presence or not of 50 µM orthovanadate. Their corresponding glucosides (i.e. scopolin, skimmin, fraxin and esculin, respectively) were analysed by HPLC. These analyses revealed that orthovanadate had no effect on the uptake of the non-catechol coumarins scopoletin and umbelliferone (Fig. 1c). However, orthovanadate almost abolished root uptake of fraxetin and esculetin (Fig. 1c and Supporting information Fig. S1). 50 µM orthovanadate reduced fraxin and esculin content by more than 90%. Importantly, no fraxetin or esculetin was detected in roots by HPLC, demonstrating that orthovanadate prevents fraxetin and esculetin uptake and not their glycosylation. The active uptake of fraxetin was confirmed using the sulfonylurea glibenclamide, another inhibitor of ATP-dependent transport (Forestier *et al*., 2003), that, similarly to orthovanadate, inhibited fraxetin uptake and had no significant effect on that of scopoletin (Fig. S2).

**Fig. 1.**
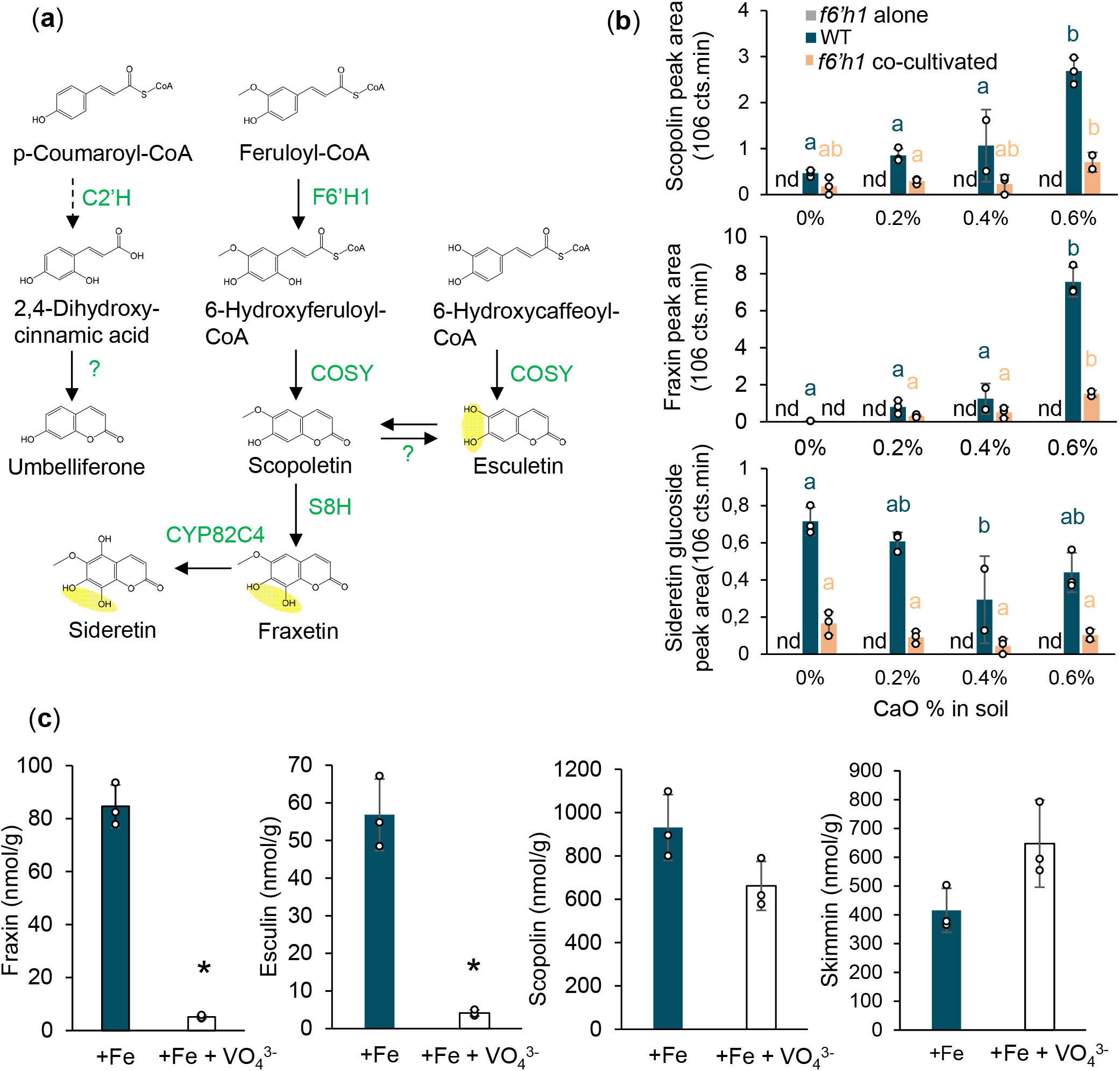
Uptake of coumarins in the presence of iron. (a) Coumarin biosynthetic pathway. Enzymes are written in capital green letters. Catechol groups are highlighted in yellow. Note that conversion of 6-Hydroxycaffeoyl-CoA into esculetin has only been demonstrated *in vitro*. Question marks indicate putative unknown enzymes. The dashed arrow represents a multi-enzymatic step where C2’H catalyses the ortho-hydroxylation of p-coumaroyl-CoA (Vialart *et al*., 2012). (b) Scopolin, fraxin and sideretin glucoside content in the roots of three-week-old *Arabidopsis thaliana* wild type (WT) and *f6’h1* mutant co-cultivated or not with the WT in soil supplemented with different CaO concentrations (%, w/w) quantified by liquid chromatography-tandem mass spectrometry (LC-MS/MS). Coumarins were identified by their retention time, MS and MS/MS profiles and quantified by integration of their aglycone counterpart EIC (extracted-ion chromatograms) signals. EIC used for the quantification of each coumarin were as follows: Scopolin : *m/z* 193.04 +/- 0.02 Sideretin glycoside : *m/z* 225.03 +/- 0.02, Fraxin : *m/z* 209.05 +/- 0.02. nd: non detected. Means within the same genotype with the same letter are not significantly different according to one-way ANOVA followed by post hoc Tukey test, P < 0.05 (n = 3 biological repeats). Bars represent means ± SD. (c) Uptake of coumarins by roots grown in hydroponic Hoagland solution containing poorly available Fe (+Fe), supplemented or not with 50 µM orthovanadate (VO_4_^3-^). Plants were kept for three days in the presence of coumarins and orthovanadate. Coumarin glucosides were analysed by HPLC. *t* test significant difference: *P < 0.05 (n = 3 biological repeats). Bars represent means ± SD.

To study the kinetics of the uptake process, a time course experiment was conducted using *f6’h1* seedlings grown on control media and transferred to a medium containing synthetic fraxetin. Samples were harvested 1, 3, 7, 24, 48 and 72 hours after transfer and fraxin accumulation was followed by HPLC (Fig. **2a**). One hour after transfer, fraxin could already be detected in the roots even though the levels were still low, indicating that both the uptake and the glycosylation, which converts fraxetin to fraxin, are fast processes. Then, the accumulation of fraxin in the roots increased regularly over the first 48 hours period of the experiment and started to decrease 72 hours after the transfer.

**Fig. 2.**
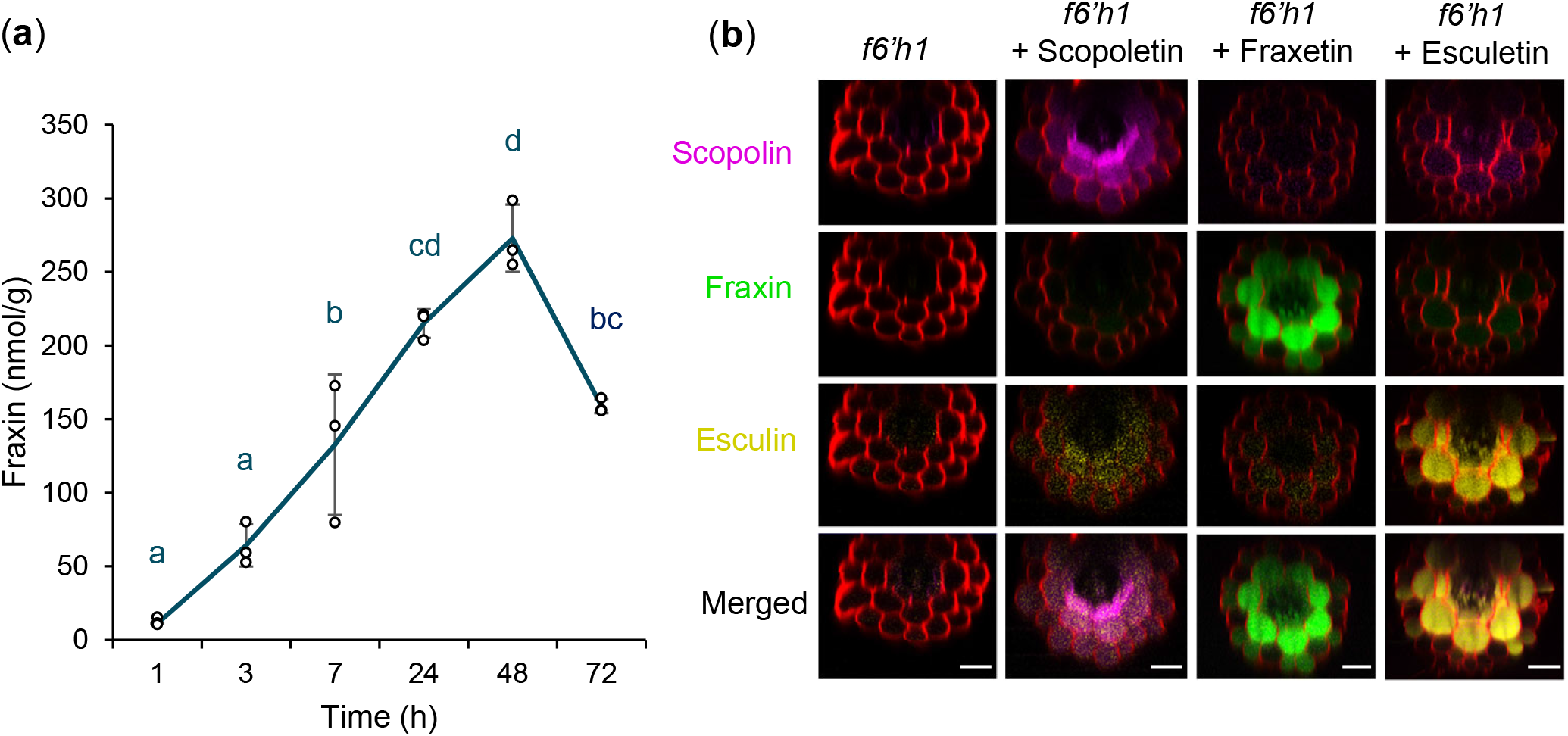
Catechol coumarins uptake kinetics. (a) Three days kinetic analysis of fraxetin uptake in *f6’h1* mutant grown in the presence of poorly available Fe and exogenous fraxetin. Time course root samples were collected and fraxin was analysed by HPLC. Means with the same letter are not significantly different according to one-way ANOVA followed by post hoc Tukey test, P < 0.05 (*n* = 3 pools of at least 50 seedlings each). Error bars show ± SD. Blue lines are used to connect the means. (b) Localisation of coumarins (i.e. scopolin: purple, fraxin: green, esculin: dark yellow) in *Arabidopsis thaliana f6’h1* mutant seedlings grown during 3 days in the presence of synthetic scopoletin, fraxetin or esculetin. Images were obtained by spectral deconvolution and are representative of observations made in three independent experiments where the roots of five to six seedlings were independently analysed. Bar = 25 µm.

Spectral imaging was used to investigate the localization of scopolin, fraxin and esculin after the uptake of their corresponding aglycone form by *f6’h1* mutants in the presence of poorly available Fe. For this purpose, *f6’h1* mutant was grown in a medium containing 25 µM of iron chloride (FeCl_3_) at pH7 and catechol or non-catechol coumarins. Spectral imaging confirmed that seedlings were able to take up coumarins in the presence of Fe, glycosylate them and store the glucosides in the vacuoles (Fig. **2b**). Similarly to the accumulation pattern observed in response to Fe deficiency (Robe *et al*., 2021a), fraxin was mostly observed in the cortex and to a lesser extent in the epidermis of the root whereas esculin and scopolin accumulated in endodermis, cortex and epidermis. The absence of fraxetin or esculetin in root hairs (Fig. **2c**) suggests that uptaken coumarins were not just stored but could also be secreted back into the rhizosphere. These results demonstrate that catechol coumarins are taken up into Arabidopsis roots by a yet unknown ATP dependent mechanism. In contrast, non-catechol coumarins seem to passively diffuse into the root, as their Log K_ow_ suggests.

### Identification and characterization of the Fe-catechol coumarins complexes

Recently, some Fe-coumarin complexes were identified and characterised by mass spectrometry (Baune *et al*., 2020). However, these experiments were done after chromatographic separation at acidic pH (the mobile phases contained formic acid), which tends to dissociate Fe-coumarin complexes (Rajniak *et al*., 2018). Another work (Schmid *et al*., 2014) reported Fe(II)-(Scopoletin)_3_ complex, even though scopoletin lacks catechol moiety that binds Fe but couldn’t identify Fe complexes with fraxetin and esculetin.

In order to investigate Fe-coumarin complexes at circumneutral to alkaline pH we prepared scopoletin, esculetin and fraxetin solutions and mixed them to FeCl_3_ in an ammonium bicarbonate buffer at pH 7. Each mixture was analysed by direct infusion in positive and negative ESI modes. In positive ESI mode, only free coumarins showed efficient ionisation and we could barely detect Fe complexes. In negative ESI mode, Fe(III) complexes with fraxetin and esculetin were identified as singly charged species (Fig. 3). Fe complexes can be easily recognised by the characteristic isotopic profile of Fe (Tsednee *et al*., 2016): the relative abundance of naturally occurring stable Fe isotopes is approximately ^54^Fe (5.8%), ^56^Fe (91.7%), ^57^Fe (2.2%) and ^58^Fe (0.3%). When fraxetin/FeCl_3_ mixtures were infused, the main species detected was the 1:2 stoichiometry complex [Fe^III^(fraxetin)_2_-4H^+^]^-^ (*m/z* 467.99), but we could also observe the 1:3 [Fe^III^(fraxetin)3-4H^+^]^-^ (*m/z* 676.03) and the 2:4 [Fe^III^_2_(fraxetin)_4_-8H^+^]^-^ (*m/z* 936.98) (Fig. **3a**). The analysis of esculetin/FeCl_3_ solutions led to the identification of [Fe^III^(esculetin)_2_-4H^+^]^-^ (*m/z* 407.96) (Fig. **3b**). No Fe complexes with scopoletin were observed. It is noteworthy that even without Fe addition small quantities of Fe complexes with fraxetin were detected (Fig. **3a**), most probably due to Fe contamination of the infusion system. The decrease of free coumarin signal intensity after Fe addition was much higher for fraxetin (90%) than for esculetin (25%) (Fig. **3a** and **3b** compared to Fig. **3c**), showing that fraxetin is a stronger Fe chelator than esculetin at circumneutral pH. Fraxetin is mostly produced in alkaline soil (Fig. **1b**), suggesting that Fe-fraxetin complexes are more stable under this condition. Interestingly, when Fe was added we could detect significant amounts of oxidised free fraxetin (fraxetin semiquinone, *m/z* 206.0221) (Fig. **3a**), while in the absence of Fe only the reduced form of fraxetin was detected (*m/z* 207.0351) (Fig. **3c**). This observation suggests that, while most of the solubilised Fe(III) forms stable chelates with fraxetin, the redox dissociation of this complex may give rise to a small fraction of soluble Fe^2+^ and free oxidised fraxetin.

**Fig. 3.**
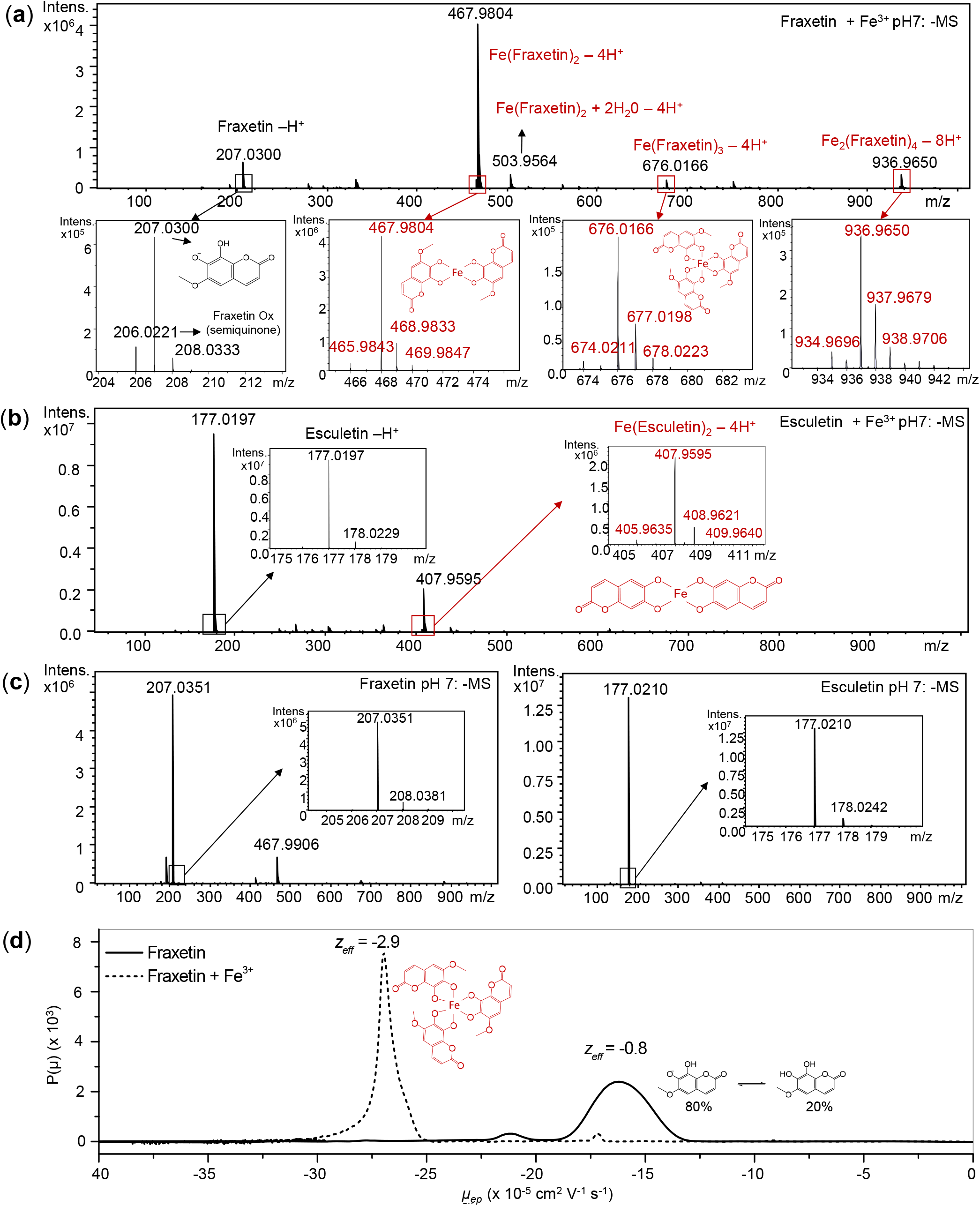
Identification of Fe-fraxetin and Fe-esculetin complexes. (a) Direct infusion ESI-QTOF MS analysis of a 2:1 (mol:mol) mixture of synthetic fraxetin and Fe(III) at pH 7. Note that [Fe(fraxetin)_2_-4H^+^]^-^ can be detected even in the absence of an external iron source, certainly due to iron contamination. (b) Direct infusion ESI-QTOF MS analysis of 2:1 (mol:mol) mixture of synthetic esculetin and Fe(III) at pH 7. (c) Direct infusion ESI-QTOF MS analysis of fraxetin (left) and esculetin (right) at pH 7 without any Fe addition. (a-c) ESI was operated in negative mode. Iron complexes were identified by their characteristic isotopic pattern and are highlighted in red. Black and red boxes show a zoom on the MS signal of the main coumarin species. (d) Electrophoretic mobility distributions for fraxetin (solid line) and a 2:1 (mol:mol) mixture of fraxetin and Fe(III) (dashed line) at pH 7 in a 50 mM Ammonium carbonate buffer. The effective charge (z_eff_) was calculated using the model described in (Oshima, 2001). The structure of the main iron-containing complex is highlighted in red. The abundance of the protonated and deprotonated forms of fraxetin was calculated from its effective charge and is in agreement with fraxetin’s p*K*_*a*_.

The stability of Fe-fraxetin complexes at different pH was assessed by mass spectrometry (Fig. S3). While no significant differences in signal intensity of the complex (*m/z* 467.99) were observed at pH 8 and 7, it slightly decreased at pH 6, 50% of the complex was lost at pH 5 and almost no complex remained at pH 3. The loss of the complex signal resulted in an increase of free fraxetin signal, thus confirming that the Fe-fraxetin complex dissociates at acidic pH (Fig. S3).

Contrary to a previous work (Baune *et al*., 2020), all the Fe-coumarin complexes detected in our study contained mainly Fe(III). Singly charged species of Fe(II) complexes would contain one more proton than Fe(III) ones and consequently are 1Da heavier. The isotopic profiles of 467.99 and 676.01 *m/z* peaks were very similar to theoretical ones (Supporting information Table S1) making very unlikely the presence of +1Da signals belonging to Fe(II) complexes.

The ESI source is an artificial environment that induces ionisation and does not reflect the real charge of the species in their physiological state. Moreover, not all the species can be ionised with the same yield. To further elucidate if fraxetin chelates Fe(III) or Fe(II), we analysed Fe-fraxetin complexes by capillary electrophoresis, which is a milder technique that keeps molecules in their native state. Fig. **3d** shows the electrophoretic mobility distributions of fraxetin and a fraxetin/FeCl_3_ mixture obtained by transforming the temporal experimental electropherogram at pH 7 into a mobility distribution (Chamieh *et al*., 2015). Fraxetin had an average electrophoretic mobility (*µ*_*ep*_) of -16.42 ⨯ 10^−5^ cm^2^ s^-1^ V^-1^, while the Fe(III)-Fraxetin complex had a value of *µ*_*ep*_ = -27.02 ⨯ 10^−5^ cm^2^ s^-1^ V^-1^ (Fig. **3d**). By using the latter *µ*_*ep*_ values and the hydrodynamic radii values of fraxetin, (*R*_*h*_ = 0.49 nm obtained by Taylor Dispersion Analysis), and of Fe(III)-Fraxetin 1:3 complex (*R*_*h*_ = 0.77 nm obtained by modelling), the effective charge of both species could be calculated and was found to be -0.8 and -2.9 for fraxetin and the Fe(III)-fraxetin complex, respectively. The fraxetin effective charge value is in accordance with the p*K*_*a*_ = 6.61 value determined in this work (Fig. S4). The effective charge value of -2.9 obtained for the Fe-fraxetin complex indicates that the main species existing at pH 7 is the 1:3 stoichiometry Fe(III)-fraxetin complex (Fig. **3d**, Fig S5 and Table S2) where one Fe^3+^ atom is coordinated by three fraxetin molecules, each having lost two protons, resulting in a global charge of -3. Other stoichiometry complexes might also be present but at very low concentrations. The higher MS intensity observed for the 467.98 *m/z* peak corresponding to the 1:2 complex may be due to a better ionisation yield of this species and/or to *in source* dissociation of the 1:3 complex.

In analogy with phytosiderophores, we wondered if coumarins could form complexes with other transition metals. We investigated the interaction of fraxetin with the divalent cations Ni^2+^, Zn^2+^, Mn^2+^ and Cu^2+^ and the trivalent cation Al^3+^. The identification of metal-coumarin complexes relied, as for Fe, on metal-specific isotopic signatures of the metal complex spectra (Table S3). We could detect low signals corresponding to fraxetin complexes with Zn, Al, Ni, Mn and Cu in a metal-ligand stoichiometry of 1:2 (Fig. S6). However, for all of them, most of the fraxetin remained as free ligand, showing that the chelation efficiency of these metals by fraxetin is very low compared to Fe. These results demonstrate that fraxetin is a strong and highly specific Fe chelator compared to other transition metals.

### Uptake of Fe-coumarin complexes can rescue *irt1* and *bhlh121* mutants defective in Fe^2+^ uptake

*f6’h1* mutants growth defects can be complemented by growing WT plants in close vicinity to them or by adding synthetic coumarins to the culture medium (Tsai *et al*., 2018; Vanholme *et al*., 2019). Nevertheless, how coumarins can complement *f6’h1* is still an open question. The stability of Fe(III)-fraxetin complexes characterised in this work supports the appealing hypothesis that catechol coumarins may enter the root as Fe-coumarin complexes, similar to phytosiderophores in grass species. To test this hypothesis, we used *irt1* and *bhlh121* mutants. The *irt1* mutant is unable to take up Fe^2+^ ions from the soil, leading to strong chlorosis of the plant (Vert *et al*., 2002). The newly described *bhlh121* mutant is unable to activate the Fe deficiency responses (Vert *et al*., 2002; Kim *et al*., 2019; Gao *et al*., 2020a). These two mutants display strong chlorosis and cannot grow correctly unless high concentration of Fe is provided (Vert *et al*., 2002; Kim *et al*., 2019; Gao *et al*., 2020a).

We first investigated the effect of different fraxetin concentrations on *bhlh121* mutant phenotype. For this purpose, *bhlh121* mutant was grown for one week on Hoagland medium containing 75 µM Fe-EDTA and then transferred to a Hoagland medium containing poorly available Fe. As expected, when *bhlh121* mutant was grown in the presence of 25 µM FeCl_3_ at pH 7, plants displayed strong chlorosis phenotype, with necrotic spots in younger leaves (Fig. **4a**). As little as 5 µM fraxetin supply to the medium partially rescued the phenotype. At this concentration, no necrotic spots were detectable even though leaves were still chlorotic. When the concentration of fraxetin was increased, the *bhlh121* chlorotic phenotype progressively disappeared, in a dose dependent manner. Optimal growth was observed for fraxetin concentrations ranging from 70 µM to 100 µM (Fig. **4a** and Fig. S7a). We also confirmed by spectral imaging that fraxetin was taken up by the root and glycosylated into fraxin in *bhlh121* mutant (Fig. S7b).

Even though *bhlh121* chlorotic phenotype was rescued by external addition of fraxetin, we could not exclude that this effect was due to the uptake of fraxetin-reduced Fe^2+^ ions by additional ferrous iron transport systems. To confirm that coumarins directly enter the root bound to Fe(III), we used the strong Fe(II) chelator ferrozine (3-(2-Pyridyl)-5,6-diphenyl-1,2,4-triazine-4′,4′′-disulfonic acid) to chelate all the free Fe^2+^ ions that could have been reduced by fraxetin. As expected, the ability of fraxetin to rescue the *bhlh121* and the *irt1* mutant Fe-deficiency phenotypes was not reduced when ferrozine was also present in the culture medium and plants displayed the same phenotype as those grown without ferrozine (Fig. **4b**). These results indicate that the complementation of *bhlh121-2* and *irt1* mutant phenotypes by fraxetin does not depend on Fe^2+^ ions and is thus most probably due to the direct uptake of Fe(III)-fraxetin complexes by plant roots.

The Fe-deficiency phenotype of *bhlh121, irt1* and other Fe acquisition mutants such as *fro2* and *f6’h1*, directly grown on Hoagland medium containing poorly available Fe, was rescued by the addition of fraxetin and to a lesser extent of esculetin (Fig. **4d-f**). In contrast, scopoletin, that lacks the catechol moiety, was unable to complement any mutant (Figures 4d-f). It is noteworthy that *irt1* produces catechol coumarins in both Fe sufficient and deficient growth conditions, contrary to *bhlh121* mutant (Fig. S8). This difference might explain why *irt1* mutant grows better than *bhlh121* when it is cultivated at pH 7 without the addition of external catechol coumarins. To better understand the effect of coumarin secretion on *irt1* mutant Fe nutrition,

**Fig. 4.**
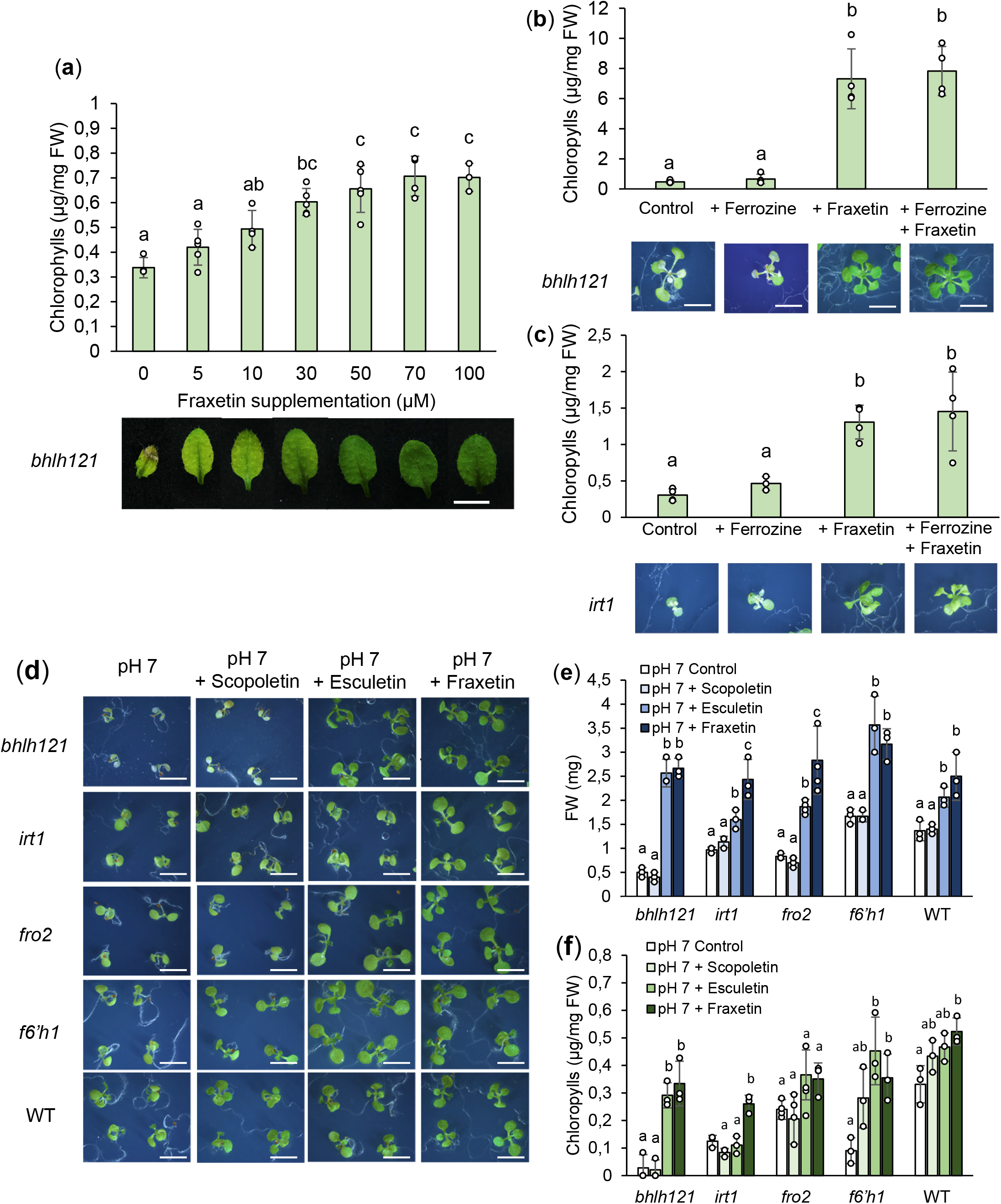
Effect of coumarins on *Arabidopsis thaliana* iron acquisition mutants. (a) Dose dependent chlorophyll content (top) and leaf phenotype (bottom) of *bhlh121* mutant supplemented with fraxetin. Fraxetin was added at different concentrations in Hoagland medium containing poorly available Fe. (b) Chlorophyll content (top) and phenotype (bottom) of *bhlh121* mutant grown in the absence of Fe^2+^ ions. The strong Fe^2+^ chelator ferrozine was added to the medium at 75µM. (c) Chlorophyll content (top) and phenotype (bottom) of *irt1* mutant grown in the absence of Fe^2+^ ions. The strong Fe^2+^ chelator ferrozine was added to the medium at 75µM. (d) Phenotype of several iron acquisition mutants grown for ten days on poorly available Fe Hoagland medium supplemented with scopoletin, esculetin or fraxetin. Plants were directly sown on coumarin containing medium. (e) Fresh weight content of the shoots of 10 day old plantlets showed in (D). (f) Chlorophyll content of the shoots of 10 day old plantlets showed in (d). (a, b, c, e and f) Means within each genotype with the same letter are not significantly different according to one-way ANOVA followed by post hoc Tukey test, P < 0.05 (*n* = 3 biological repeats). Bars represent means ± SD.

we investigated how daily renewing of the hydroponics culture medium affected *irt1* phenotype. *Irt1* mutant remained chlorotic when the medium was daily changed whereas this symptom disappeared when the medium was kept for two weeks (Fig. S9). This later experiment strongly suggests that the accumulation of secreted coumarins in culture medium improves *irt1* Fe nutrition.

Since ferritin accumulation is a proxy of Fe content in plants, we performed Western Blot on *irt1* seedlings grown on media containing poorly available Fe and different coumarins. *irt1* mutant accumulated very low levels of ferritins in leaves. Ferritins accumulation increased when *irt1* mutant was grown with fraxetin but not in the presence of scopoletin or esculetin (Fig. S10). Since esculetin could only partially complement the phenotype of *irt1* and *bhlh121* mutants and its uptake did not result in ferritins accumulation, it is likely that the main role of this coumarin is not Fe transport into the roots. Moreover, the secretion of esculetin is rather limited (Ziegler *et al*., 2017; Tsai *et al*., 2018; Terés *et al*., 2019) and its specific role in plant Fe nutrition, if any, is still to be determined.

### Fraxetin uptake increased the iron content in plant tissues

To confirm that fraxetin is selectively taken up by the root in the form of Fe(III)-fraxetin complexes, we measured the metal content of leaves and roots of *bhlh121* and *irt1* mutants grown with fraxetin supplementation. The analysis revealed that Fe content increased in both roots and shoots of *irt1* and *bhlh121* mutants grown with fraxetin (Fig. **5**). Remarkably, the Fe content increase due to fraxetin was totally abolished when orthovanadate was also present in the culture media. Since we have shown that fraxetin uptake is also abolished by orthovanadate (Fig. **1c**), these results demonstrate that a yet unknown ATP dependent transport system takes up Fe and fraxetin jointly. It is noteworthy that the application of fraxetin did not result in an increase of other, non-ferric metals.

**Fig. 5.**
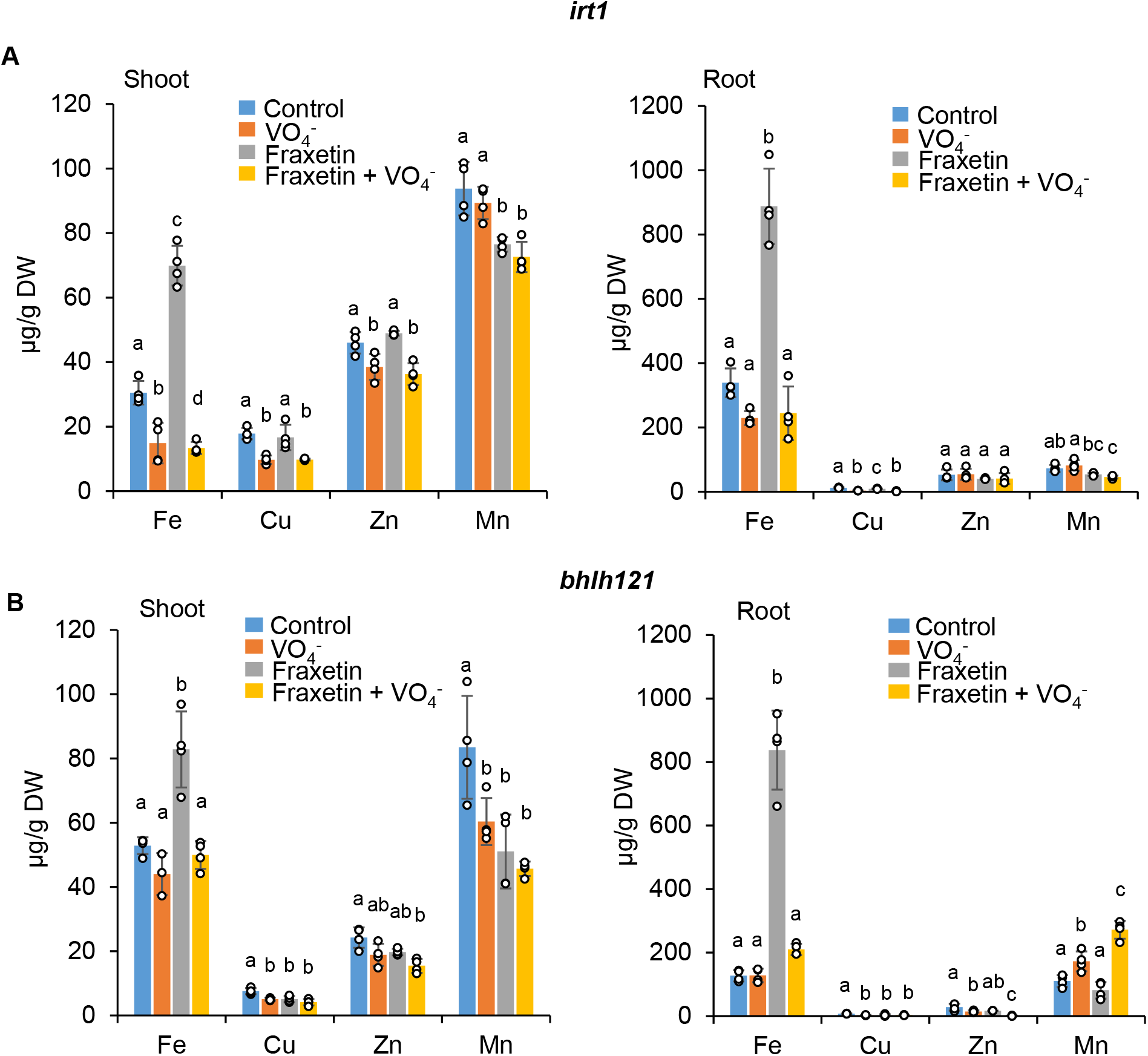
Effect of fraxetin treatment on the accumulation of iron in *Arabidopsis thaliana* iron acquisition mutants. (a) Shoots and (b) roots iron and transition metals content of *irt1* and *bhlh121* mutants grown for 10 days with fraxetin in the presence or not of 50 µM orthovanadate. *irt1* and *bhlh121* mutants were grown for 10 days in control agar plates and then transferred for 10 days to agar plates containing poorly available Fe supplemented with 100 µM fraxetin. Means within the same metal with the same letter are not significantly different according to one-way ANOVA followed by post hoc Tukey test, P < 0.05 (*n* = 4 biological repeats). Bars represent means ± SD. Fe: iron, Cu: copper, Zn: zinc and Mn: manganese.

### Uptake of Fe(III)-fraxetin is not restricted to Arabidopsis

Several eudicots such as tomato (*Solanum lycopersicum*) are able to take up fraxetin into the root (Robe *et al*., 2021a). To test if the uptake of Fe-coumarin complexes observed for Arabidopsis can also rescue Fe acquisition mutants in other Strategy I plants, we grew the T3238*fer* tomato mutant, unable to activate the Fe deficiency response, with or without fraxetin (Ling *et al*., 2002). After three weeks of growth in Hoagland medium containing 75 µM Fe-EDTA, plants were transferred to a pH 7.5 medium containing 25 µM FeCl3 supplemented or not with 100 µM fraxetin. One week after transfer in alkaline conditions, T3238*fer* mutants were chlorotic. However, T3238*fer* mutants grown with fraxetin did not display chlorotic phenotype (Fig. S11), strongly suggesting that Fe(III)-fraxetin complexes are also taken up in tomato. Additionally, HPLC analysis of root extracts showed that the T3238*fer* mutant can glycosylate the uptaken fraxetin, converting it to fraxin (Fig. S11). This observation indicates that fraxetin can prevent chlorosis in other eudicots than *Arabidopsis thaliana*. The existence of S8H orthologues in several eudicot species strongly suggests that the uptake of Fe(III)-fraxetin complexes in eudicot plants is a widespread strategy for Fe acquisition (Rajniak *et al*., 2018).

## DISCUSSION

### Uptake of Fe(III)-coumarin complexes as an alternative iron uptake system

Coumarins recently emerged as key metabolites for iron acquisition in non-grass species (Fourcroy *et al*., 2014; Schmidt *et al*., 2014; Rajniak *et al*., 2018; Tsai *et al*., 2018); however, the mechanisms by which these compounds improve iron nutrition were still unclear. Several studies show that catechol coumarins are involved in Fe reduction and/or Fe chelation (Fourcroy *et al*., 2016; Sisó-Terraza *et al*., 2016; Rajniak *et al*., 2018; Tsai *et al*., 2018). We recently showed that the catechol coumarin fraxetin is taken up by several non-grass species, suggesting that it might be a widespread phenomenon occuring within the plant kingdom (Robe *et al*., 2021a). Whether this uptake is accompanied by an uptake of Fe was still an open question (Rodríguez-Celma & Schmidt, 2013; Tsai *et al*., 2018). By complementing mutants unable to take up Fe(II) (i.e. *irt1* and *bhlh121*) with synthetic fraxetin, we demonstrated that Arabidopsis has the ability to directly take up Fe(III) from the growth media. Complementation assays conducted in the presence of ferrozine confirmed this observation. In addition, we showed that at pH 7 fraxetin and Fe can interact *in vitro* and form highly specific Fe(III)-fraxetin complexes with a 1:3 metal:ligand stoichiometry, similar to those observed for phytosiderophores. Since both fraxetin and fraxetin-mediated Fe uptake are abolished in the presence of orthovanadate, our results strongly suggest that fraxetin participates to the plant Fe nutrition by forming stable chelates with Fe(III) that are directly taken up by the plant root system (Fig. **6**) through a specific ATP dependent transporter. Fe(II) is not necessarily released from fraxetin by reductive chelate splitting. The log K_ow_ value of aglycone catechol coumarins (around 1.5) suggests a passive diffusion through the epidermis. However, the reduced uptake of fraxetin and esculetin observed in roots of plants treated with vanadate or glibenclamide clearly demonstrates that the uptake of catechol coumarins is ATP dependent.

**Fig. 6.**
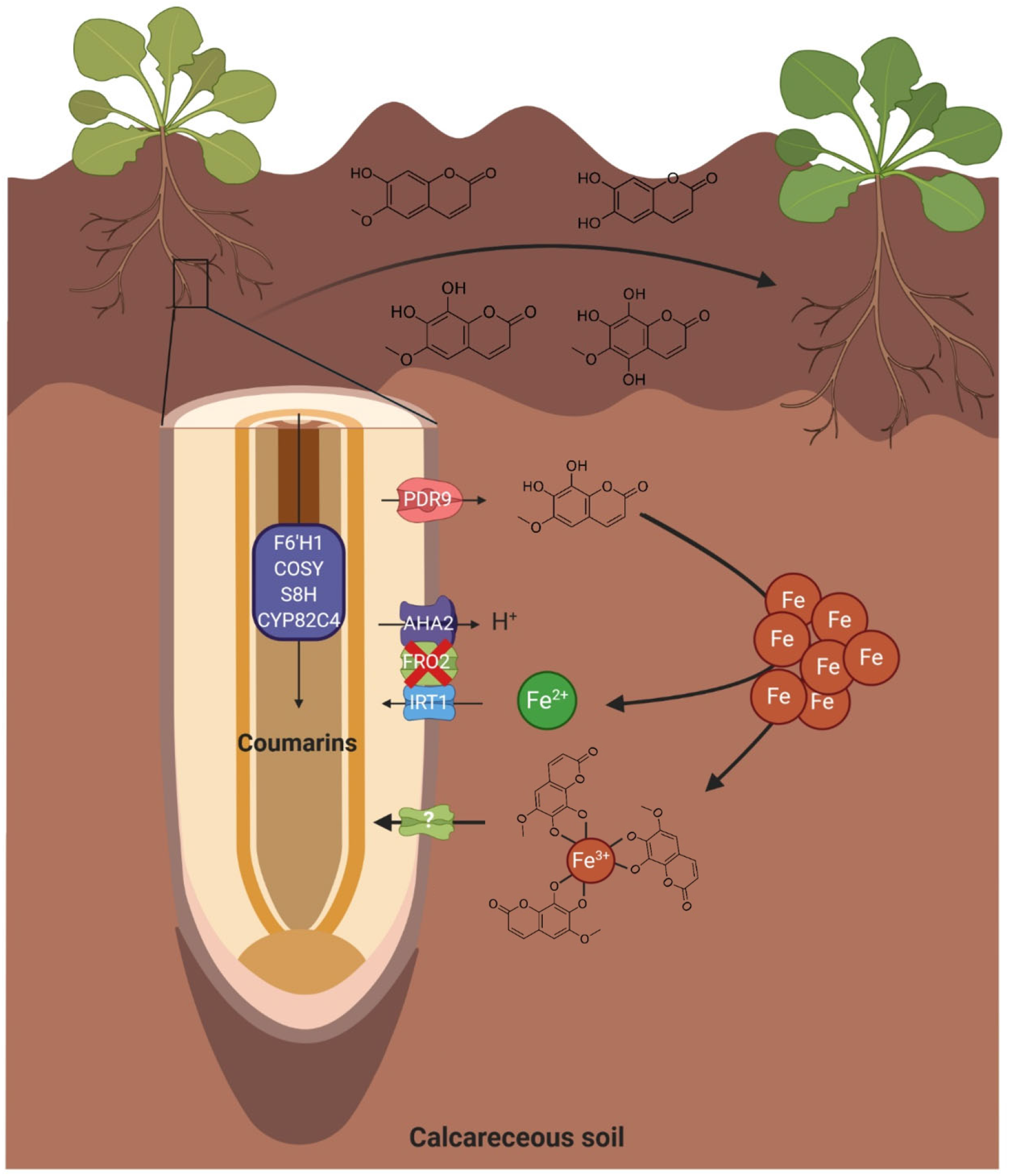
Proposed model for Fe-coumarins uptake by *Arabidopsis thaliana* roots. In calcareous or alkaline soils, FRO2/IRT1 high affinity Fe^2+^ transport system is almost inactive as FRO2 activity is strongly inhibited at high pH. As a response to this growth condition, *Arabidopsis thaliana* produces and secretes several coumarins such as scopoletin, esculetin, fraxetin and sideretin. Once secreted by PDR9, catechol coumarins will solubilise iron hydroxides by forming stable complexes with Fe(III) and also by reducing Fe(III) to Fe(II). Reduced Fe will be taken up by IRT1 and Fe(III)-coumarin complexes will be taken up by a yet unknown transporter localised at the plasma membrane. The main catechol coumarin involved in Fe(III) uptake is fraxetin and mostly act in a 3 fraxetin to 1 Fe ratio.

It was previously shown that coumarins were unable to complement *fro2* and *irt1* mutants (Fourcroy *et al*., 2016). However, the major role of pH in coumarin biosynthesis and secretion was not well understood at that time and the coumarin-containing culture media used for these experiments were obtained by cultivating plants in the absence of Fe at acidic pH. Fraxetin was certainly not produced in those conditions, which explains why *fro2* and *irt1* could not be complemented.

Here, we provide evidence of a long time suspected parallel Fe acquisition system in non-grass species. The inhibitory effect of orthovanadate treatment on catechol coumarin uptake could be due to the inactivation of plasma membrane localised ATPases, such as AHA2, resulting in a hampered proton gradient. This would suggest that a co-transport of Fe-coumarin complexes with protons is occuring. In analogy with the uptake of phytosiderophores, an YSL transporter might also be involved in the uptake of Fe-coumarin complexes. YSL transporters appear as good candidates since ZmYS1 is involved in iron bound siderophore uptake and the peanut AhYSL1 is responsible for siderophore uptake (Xiong *et al*., 2013). Nevertheless, it cannot be excluded that the uptake of catechol coumarins relies on the activity of an orthovanadate sensitive transporter, such as the ATP-binding cassette transporters (ABC) types, to which PDR9 belongs. Extensive studies will be necessary to identify and characterise the transporter involved in the uptake of Fe-coumarin complexes.

### The advantage of a parallel iron uptake system for strategy I plants

From an ecological point of view, the existence of a chelation-based Fe(III) acquisition system for Strategy I plants might be a strong advantage when soil conditions, such as high pH (Römheld & Marschner, 1983; Terés *et al*., 2019), prevent Fe reduction by FRO2, necessary for the uptake of this nutrient through the high affinity Fe(II) transporter IRT1. It may also explain how non-grass species, including Arabidopsis, can thrive in carbonate-rich soils with low bioavailability of Fe. For instance, it was shown that fraxetin secretion of 22 different Arabidopsis accessions is highly correlated with plant Fe content and that accessions abundantly secreting fraxetin have the best growth parameters (Tsai *et al*., 2018). In addition, it was reported that a high fraxin content in roots and a high capacity for fraxetin secretion are responsible for the growth improvement of local Arabidopsis populations adapted to alkaline soils (pH 7.5) (Terés *et al*., 2019). On the contrary, root ferric reductase activity is similar between this population and the pH sensitive ones (Terés *et al*., 2019).

The identification of a second Fe acquisition system is a major step towards the complete understanding of Fe uptake in non-grass plant species and might have important implications for improving cultural practices to prevent Fe deficiency. The selection of plants being able to secrete higher amounts of fraxetin might significantly reduce Fe induced chlorosis in the field without adding synthetic Fe chelates and thus allow the culture and yield improvement in soils with poor Fe availability. Previous studies have demonstrated that grass species can improve Fe nutrition of non-grass plants in intercropping systems (Brown *et al*., 1991; Hopkins *et al*., 1992; Kamal *et al*., 2000; Gunes *et al*., 2007; Xiong *et al*., 2013). Hence, it can be hypothesised that the transporter involved in Fe-coumarin complexes uptake might also be involved in the uptake of phytosiderophores. However, a previous study indicates that the reverse is probably not occurring (Robe *et al*., 2021a).

## ACKOWLEDGEMENTS

We thank Sandrine Chay (BPMP, SAME platform, Montpellier, France) for technical support for plant Fe determination. We thank Dr. Marie Lopez (IBMM, Montpellier, France) for help with coumarin chemistry. We also thank Dr Brigitte Touraine (BPMP) for help with HPLC analysis of coumarins. This work was supported by the Agence Nationale pour la Recherche (ANR-17-CE20-0008-01 to CD). KR was supported by a fellowship from the Agence Nationale pour la Recherche (ANR-17-CE20-0008-01) and the BAP department of the INRAE.

## AUTHOR CONTRIBUTIONS

KR, JC, CD and EI conceived and designed the experiments. KR, MS, JC, PG and EI performed the experiments. KR, MS, JC, SH, CD and EI analysed the data. SH and VS provided access to mass spectrometry facilities. KR, JC, CD and EI wrote the paper with the help of all the authors.

## DATA AVAILABILITY

All data generated during and/or analysed during the current study are available from the corresponding author on reasonable request.

## SUPPORTING INFORMATION

Additional Supporting Information may be found in the Supporting Information section at the end of the article.

**Fig. S1** Inhibition of esculetin uptake by orthovanadate addition.

**Fig. S2** Inhibition of fraxetin uptake by the ATP-dependent transport inhibitor glibenclamide.

**Fig. S3** Analysis of the stability of Fe-fraxetin compleses at different pH.

**Fig. S4** Determination of the p*K*_*a*_ of fraxetin by capillary electrophoresis.

**Fig. S5** Proposed structures of Fe-Fraxetin complexes.

**Fig. S6** Identification of non-ferric metals and iron complexes with fraxetin.

**Fig. S7** Phenotypic effect of fraxetin application on *bhlh121* mutant.

**Fig. S8** Coumarin accumulation in roots of iron acquisiton mutants.

**Fig. S9** *irt1* mutant grown in hydroponics.

**Fig. S10** Western Blot of ferritins accumulation in *irt1* mutant supplemented or not with coumarins.

**Fig. S11** Effect of fraxetin on tomato *T3238fer* mutant.

**Fig. S12** Example of graphical determination of zeta potential (*ζ*).

**Table S1** Experimental and theoretical mass-to-charge ratios (m/z) and isotope abundance of 2:1 and 3:1 fraxetin-Fe complexes detected by Direct infusion ESI-QTOF MS analysis

**Table S2** Charge, hydrodynamic radius and electrophoretic mobility of Fe:Fraxetin complexes.

**Table S3** Natural isotope abundances of the analyzed metals

**Methods S1** Detailed protocols for determination of the pKa of Fraxetin and the Effective charge of fraxetin and Fe(III)-fraxetin complex

