## Supporting Information for "Uptake of Fe-fraxetin complexes, an IRT1 independent strategy for iron acquisition in *Arabidopsis thaliana*"

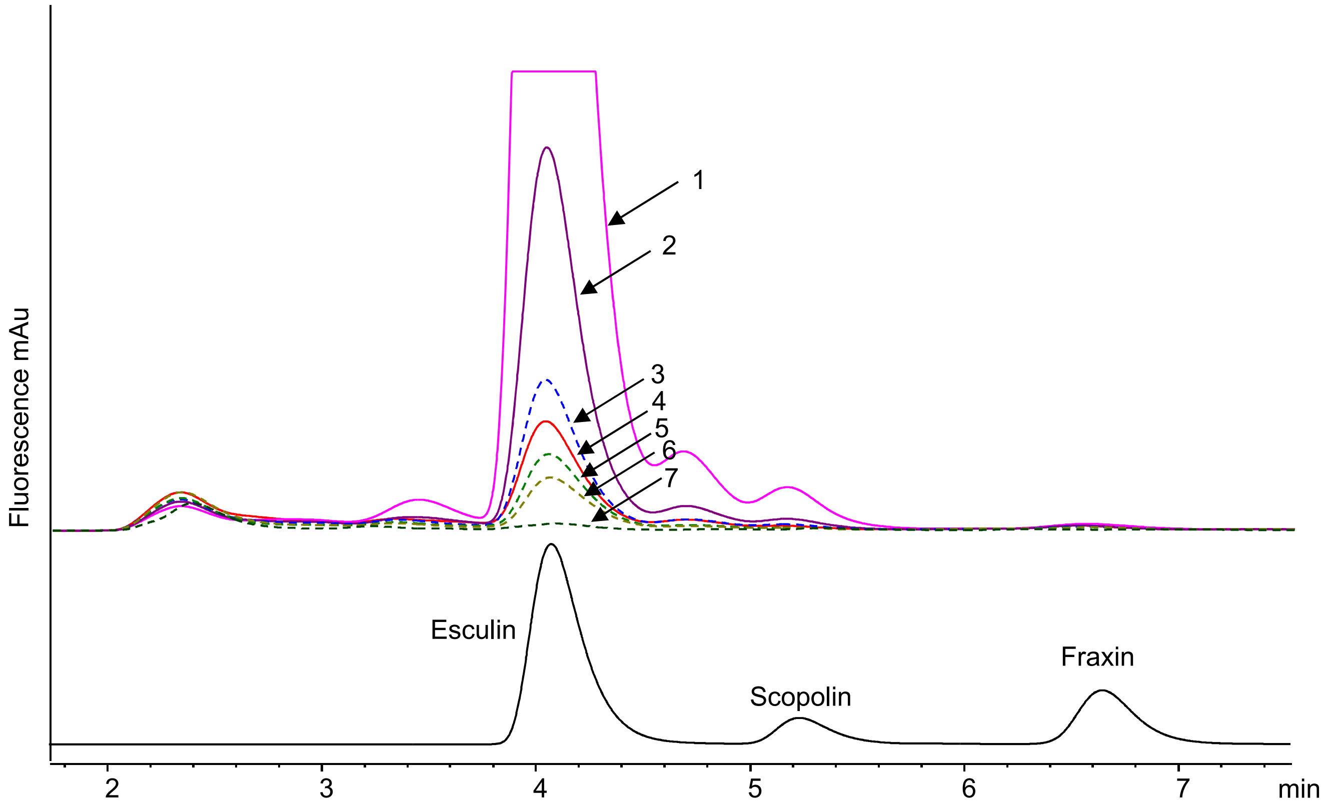


**Figure S1 : Inhibition of esculetin uptake by orthovanadate addition.** Representative fluorescence chromatograms obtained using $\lambda_{exc}$ 365 and $\lambda_{em}$ 460 nm for root extracts of Arabidopsis *f6'h1* mutant seedlings grown for 7 days on Hoagland agar plates and then transferred 7 days on Hoagland agar containing poorly available Fe and supplemented with 50µM esculetin and different concentrations of orthovanadate ($VO_4^{3-}$).
1 : + 0 µM $VO_4^{3-}$ , 2 : + 50µM $VO_4^{3-}$, 3: + 100 µM $VO_4^{3-}$, 4 : + 200 µM $VO_4^{3-}$, 5 : + 250 µM $VO_4^{3-}$, 6: + 500 µM $VO_4^{3-}$, 7: + 1 mM $VO_4^{3-}$.

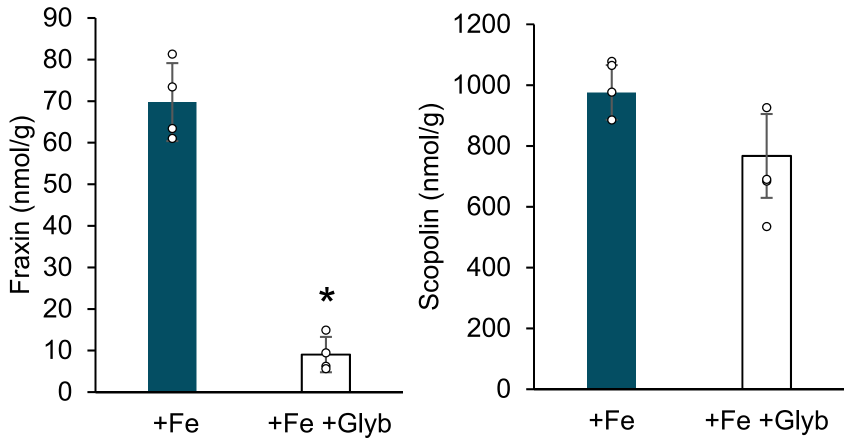


**Figure S2. Inhibition of fraxetin uptake by the ATP-dependent transport inhibitor glibenclamide.** Uptake of coumarins by roots grown in hydroponic Hoagland solution containing poorly available Fe (+Fe), supplemented or not with 150 µM glibenclamide (Glyb). Plants were kept for three days in the presence of coumarins and glibenclamide and coumarin glucosides were analyzed by HPLC. *t* test significant difference: *P < 0.05 (*n* = 4 biological repeats). Bars represent means ± SD.

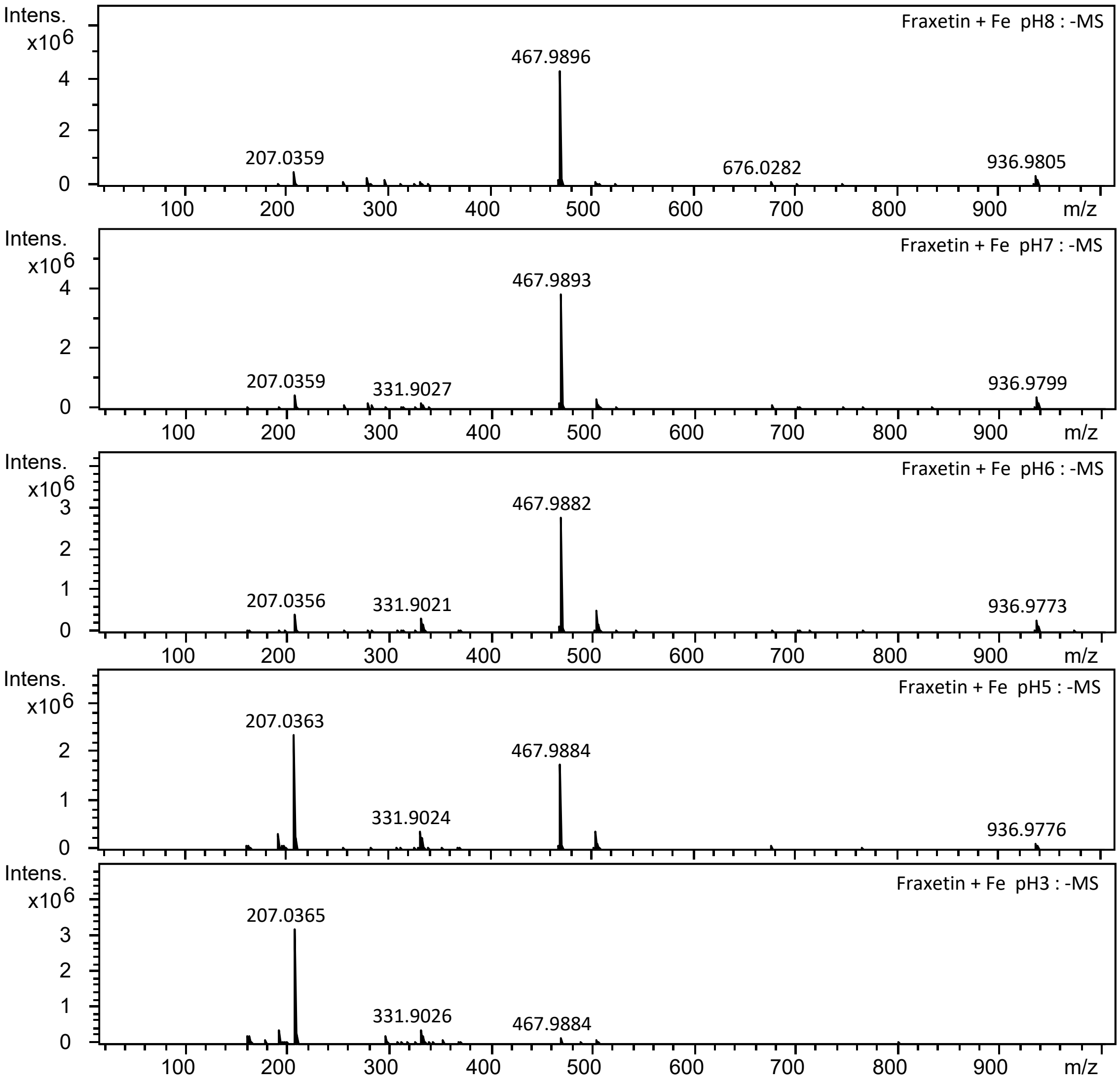


**Figure S3. Analysis of the stability of Fe-fraxetin compleses at different pH.** Direct infusion ESI-QTOF MS analysis. Stability of the complexes decreases when pH becomes more acidic. Note that free fraxetin signal increases in acidic conditions, confirming the dissociation of the complexes.

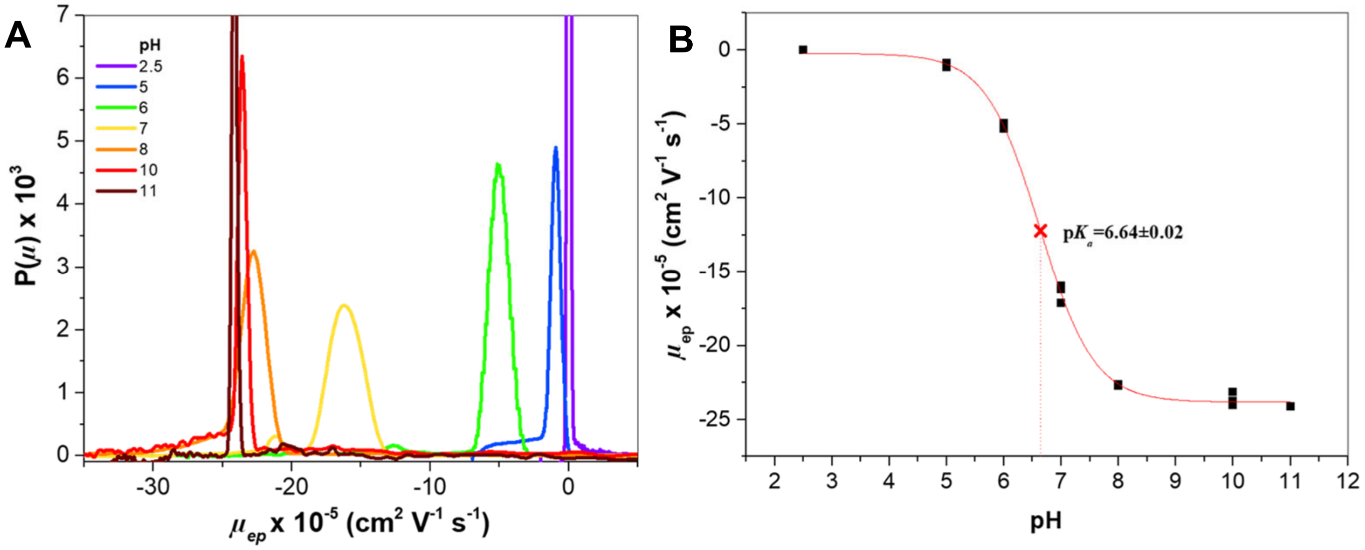


**Figure S4. Determination of the p$K_a$ of fraxetin by capillary electrophoresis.**
(a) Electrophoretic mobility distributions for fraxetin as a function of pH (n=3). (b) Evolution of the effective mobility of fraxetin as function of pH allowing the estimation of the p$K_a$ using equation (1) in the main text. Red cross represents the graphical determination of p*Ka*

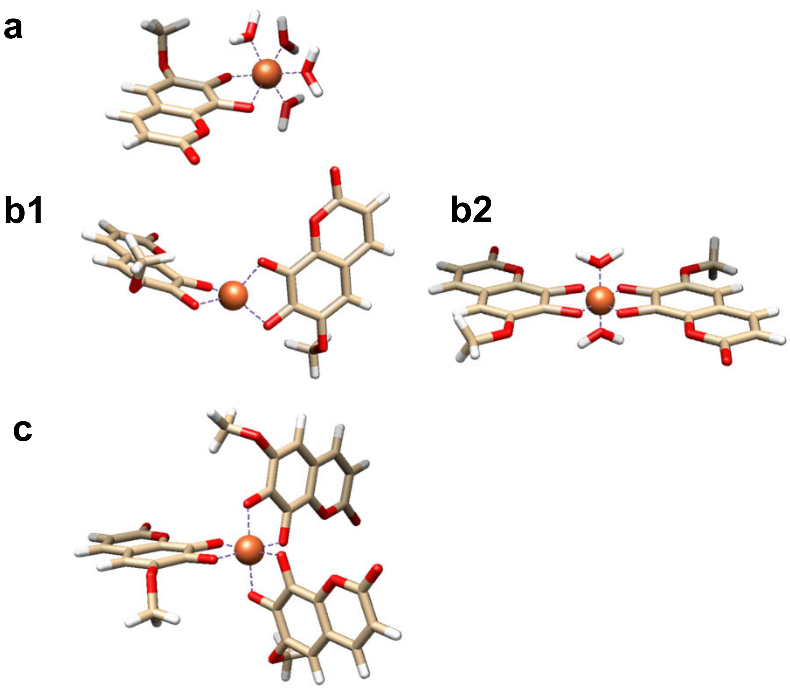


**Figure S5. Proposed structures of Fe-Fraxetin complexes. a**: Fe:Fraxetin 1:1 complex, **b1**: Fe:Fraxetin 1:2 tetrahedral complex, **b2**: Fe:Fraxetin 1:2 octahedral complex; **c**: Fe:Fraxetin 1:3 octahedral complex. When needed the coordination sphere was completed with water molecules.

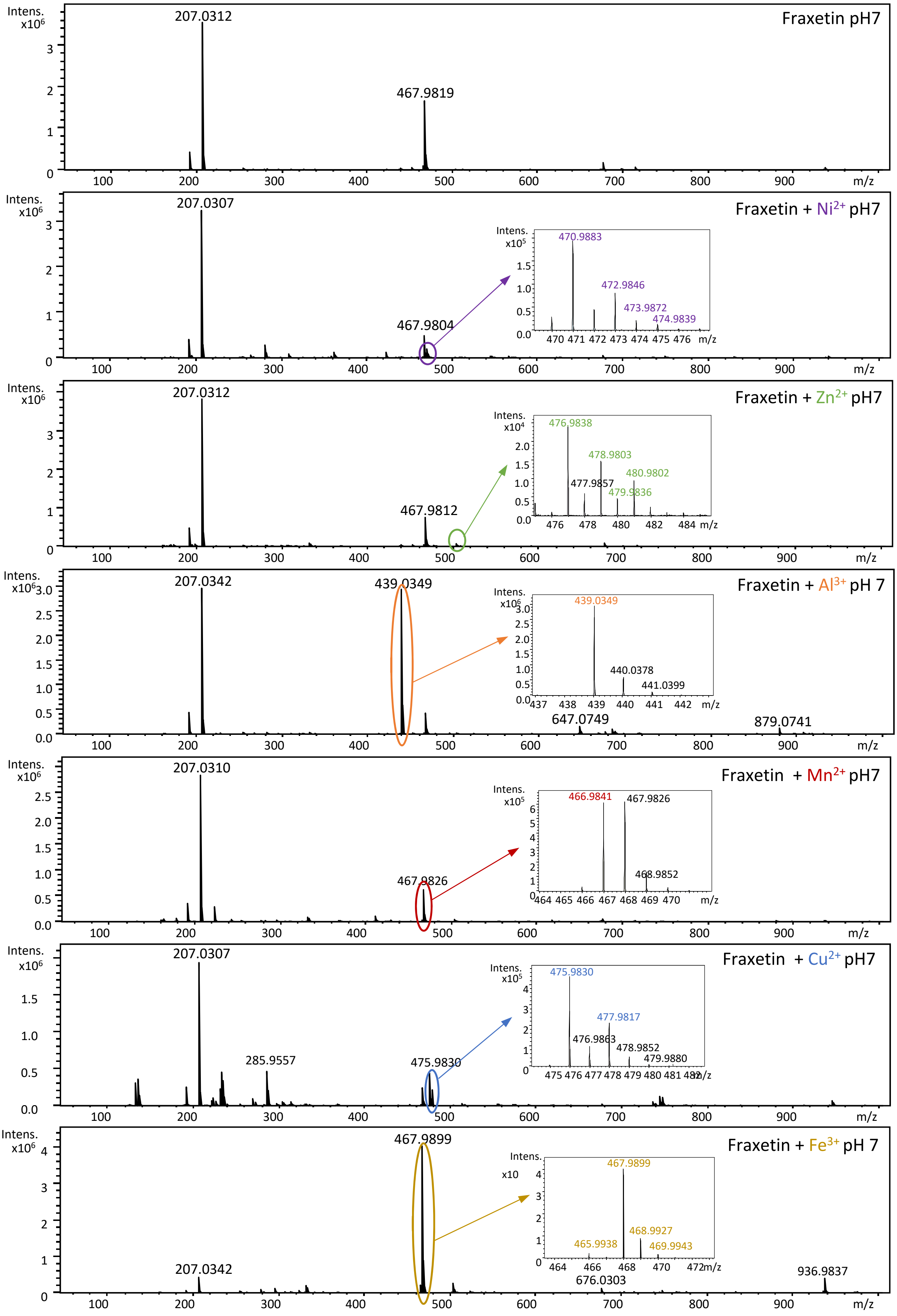


**Figure S6. Identification of non-ferric metals and iron complexes with fraxetin.** Direct infusion ESI-QTOF MS analysis of a mixture of individual transition metals and synthetic fraxetin at pH 7 are presented. All tested metal solutions were prepared from chloride salts (ie: $NiCl_2$, $AlCl_3$, $MnCl_2$, $ZnCl_2$, $CuCl_2$ and $FeCl_3$). Color ellipses show a zoom of the MS signal of the formed metal-fraxetin complexes. Isotopic signatures of each metal are highlighted in colour. All spectra were obtained in negative ESI mode with Q-TOF Maxis Impact HD (Bruker Daltonics).

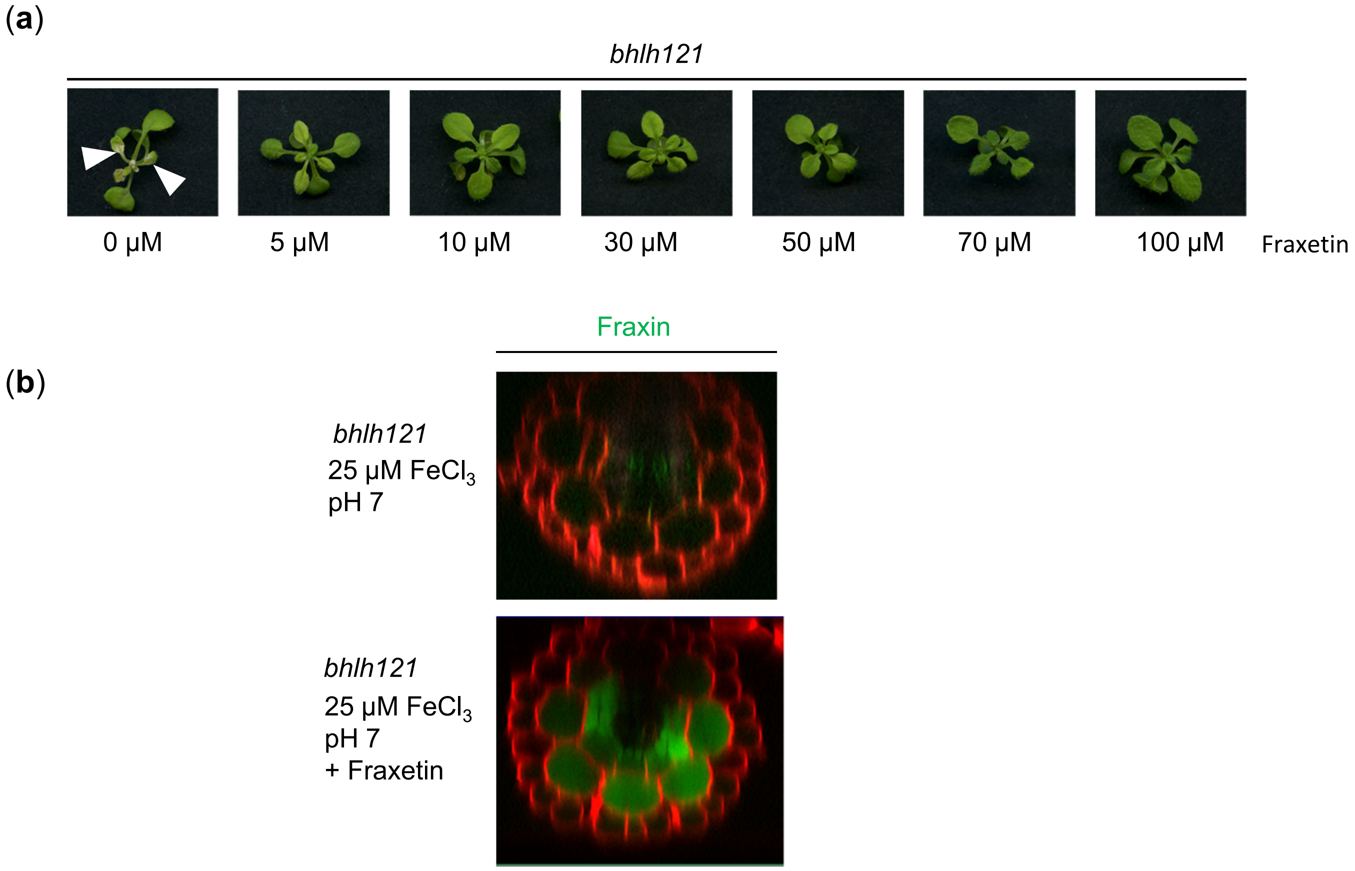


**Figure S7. Phenotypic effect of fraxetin application on *bhlh121* mutant**

(a) Dose dependent phenotype complementation of *bhlh121* mutant grown with fraxetin. Fraxetin was added at different concentrations in Hoagland medium containing 25 μM $FeCl_3$ at pH 7. (b) Spectral deconvolution image of fraxin (green) and PI (red) in *bhlh121* mutant grown in the presence of fraxetin.

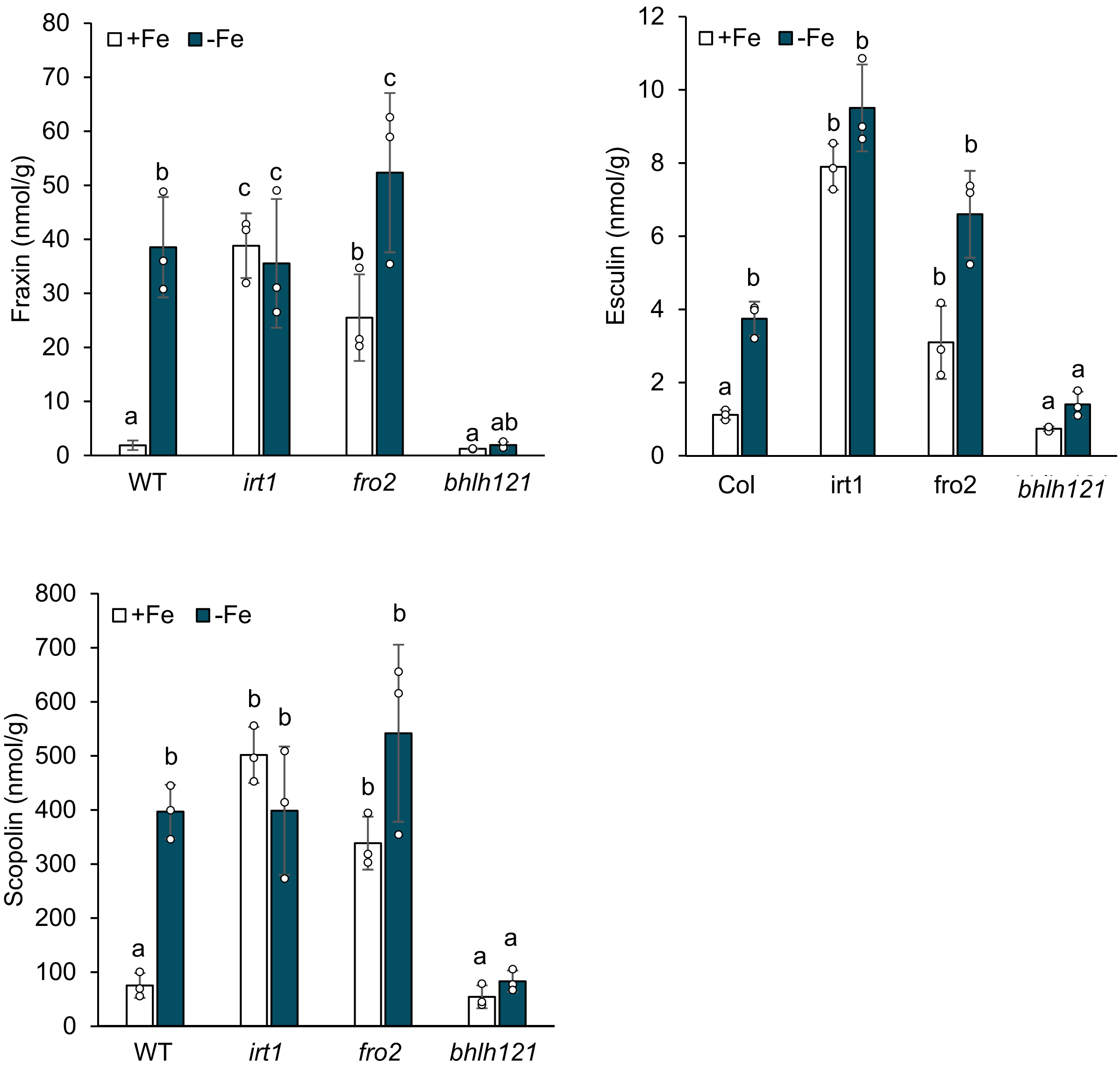


**Figure S8. Coumarin accumulation in roots of iron acquisiton mutants.** Analysis of fraxin, esculin and scopolin in roots of plants grown in iron sufficient conditions (50 µM Fe-EDTA, pH 5.5) and iron deficient conditions (0 µM Fe, pH 7). Means with the same letter are not significantly different according to one-way ANOVA followed by post hoc Tukey test, $P < 0.05$ ($n = 3$ biological repeats). Bars represent means ± SD.

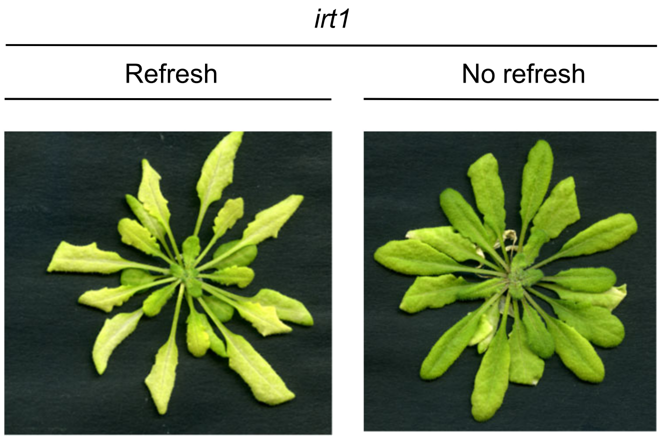


**Figure S9. *irt1* mutant grown in hydroponics.** Daily medium change effect on *irt1* mutant phenotype.

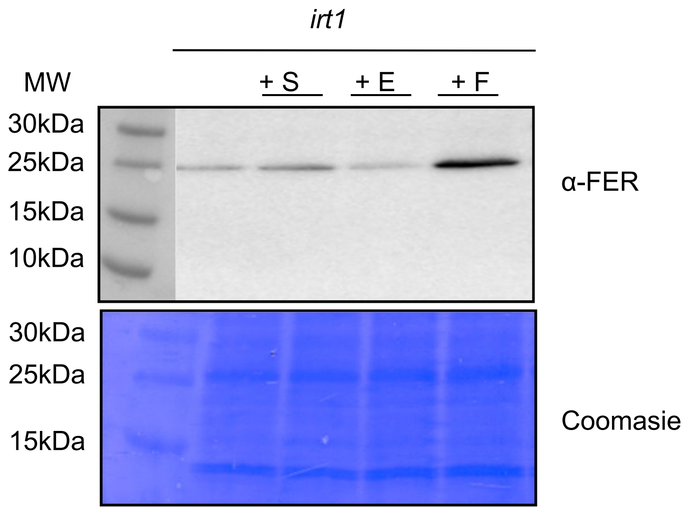


**Figure S10. Western Blot of ferritins accumulation in *irt1* mutant supplemented or not with coumarins.** Ferritins accumulation in shoots of 10-day-old *irt1* mutants grown on medium containing 25 µM $FeCl_3$ and supplemented with scopoletin (+S), esculetin (+E) or fraxetin (+F). Each loaded sample was obtained by pooling approximately 10 seedling shoots. Coomasie is shown as loading control.

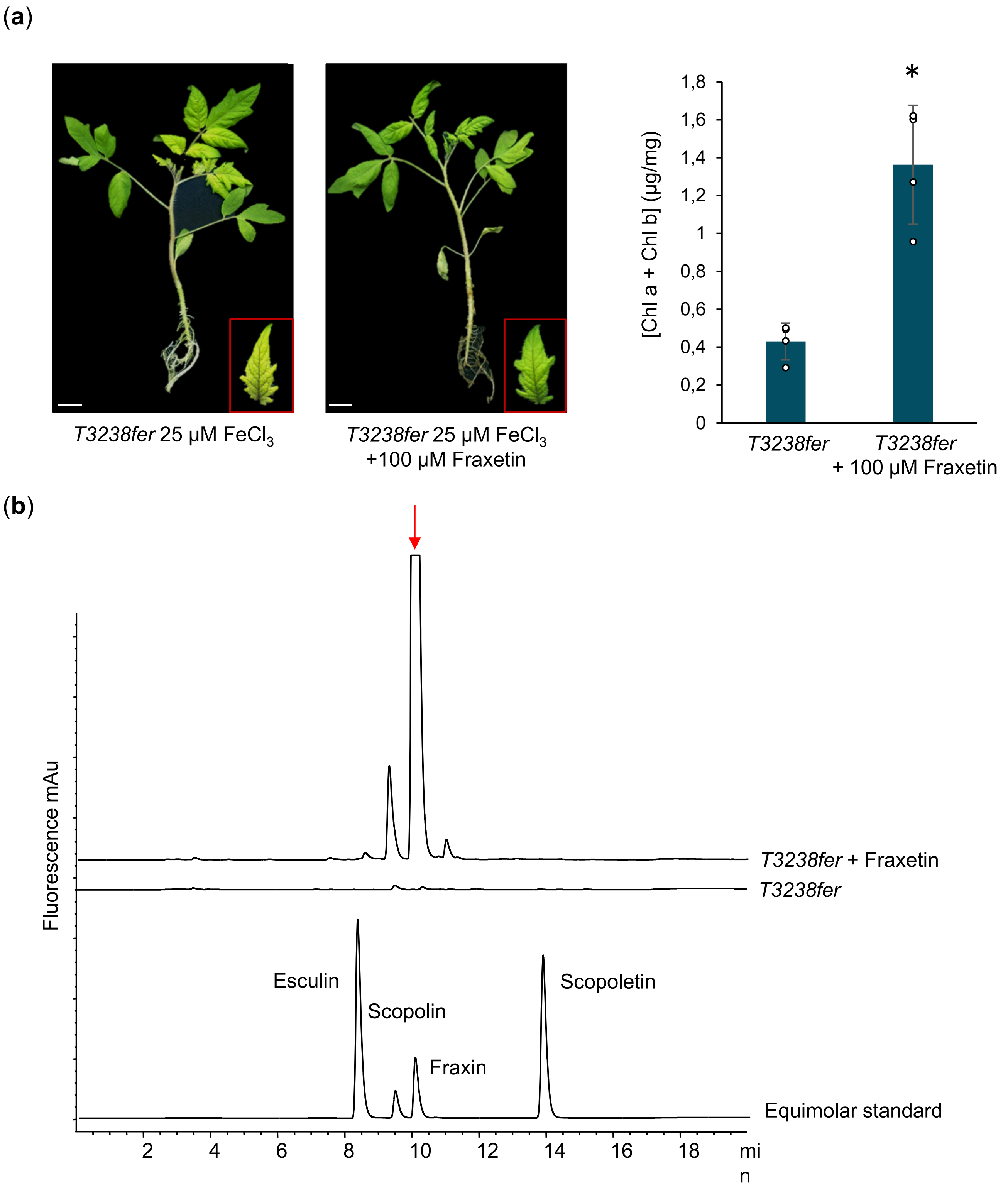


**Figure S11. Effect of fraxetin on tomato *T3238fer* mutant.**
(a) Complementation of *T3238fer* mutant with fraxetin. *T3238fer* mutant was transferred to Hoagland medium containing 25 µM $FeCl_3$ at pH 7 and supplemented or not with 100 µM Fraxetin for 1 week. Central leaves were used for chlorophyll measures. Bars represent means ± SD (n = 4). (b) Representative fluorescence chromatograms obtained using $\lambda_{exc}$ 365 and $\lambda_{em}$ 460 nm for root extracts of *T3238fer* mutant grown 2 weeks on Hoagland medium (pH 5.5) and then transferred for 7 days in Hoagland medium adjusted to pH 7.5 with KOH and supplemented or not with 100 µM fraxetin. The red arrow shows fraxin peak.

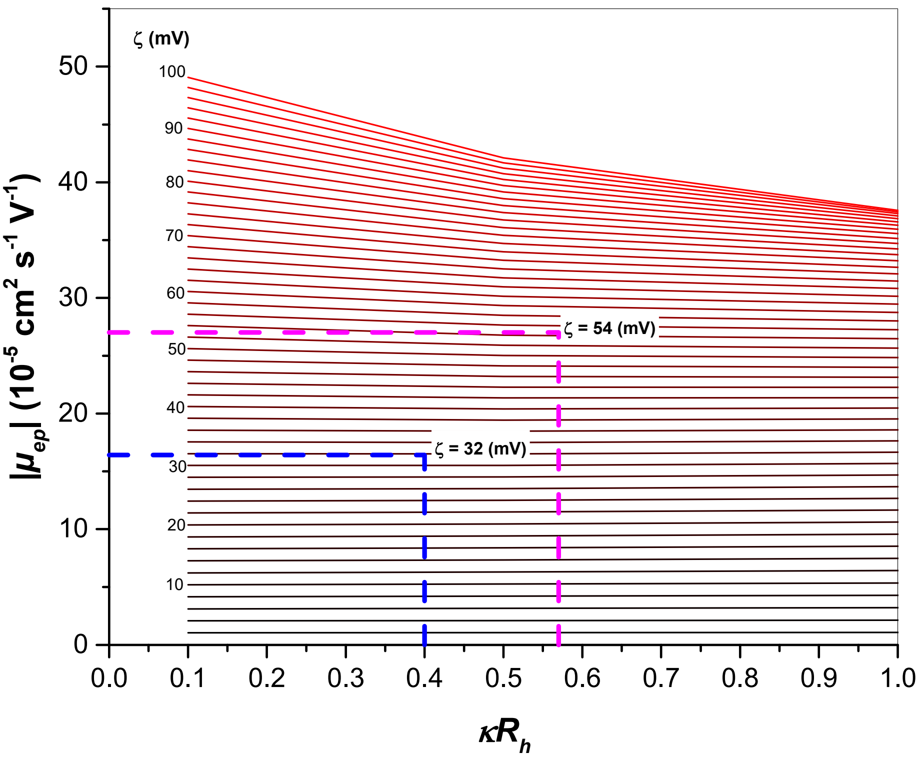


**Figure S12. Example of graphical determination of zeta potential ($\zeta$).** $\zeta$ is obtained from the experimental effective mobility (*μep*) of a solute and depending on its hydrodynamic radius and on the ionic strength of the medium according to O'Brien-White-Ohshima model (OWO) (equation (2)), in a symmetrical electrolyte of $NH_4HCO_3$ ($m+$ = 0.189, $m-$ = 0.288). The dashed lines represent the graphical determination of $\zeta$ in a 50 mM $NH_4HCO_3$ buffer pH 7 for: Fraxetin (blue dashed lines, $\kappa R_h$ = 0.39, $R_h$ = 0.49 nm, $|\mu_{ep}|$ = 16.42 × $10^{-5}$ cm$^2$ s$^{-1}$ V$^{-1}$) and Fraxetin-Fe(III) complex (3:1) (magenta dashed lines, $\kappa R_h$ = 0.57, $R_h$ = 0.77 nm, $|\mu_{ep}|$ = 27.02 × $10^{-5}$ cm$^2$ s$^{-1}$ V$^{-1}$)

| Complex | Elemental composition | Theoretical m/z | Experimental m/z | Theoretical intensity (%) | Experimental intensity (%) | Δ intensity (%) |
|---|---|---|---|---|---|---|
| $[Fe(fraxetin)_2\text{-}4H^+]^-$ | $C_{20}H_{12}O_{10}Fe$ | 465.9832 | 465.9843 | 6.4 | 5.2 | 1.2 |
| | | 466.9866 | 466.9878 | 1.4 | 1.1 | 0.3 |
| | | 467.9786 | 467.9804 | 100 | 100 | 0 |
| | | 468.9816 | 468.9833 | 24.4 | 20.5 | 3.9 |
| | | 469.9835 | 469.9847 | 5.2 | 4.1 | 1.1 |
| $[Fe(fraxetin)_3\text{-}4H^+]^-$ | $C_{30}H_{20}O_{15}Fe$ | 674.0204 | 674.0211 | 6.3 | 6.2 | 0.1 |
| | | 675.0237 | 675.018 | 2.1 | 4.2 | -2.1 |
| | | 676.0158 | 676.0166 | 100 | 100 | 0 |
| | | 677.0189 | 677.0198 | 35.5 | 35.4 | 0.1 |
| | | 678.0211 | 678.0223 | 9.5 | 8.3 | 1.2 |

**Table S1. Experimental and theoretical mass-to-charge ratios (*m/z*) and isotope abundance of 2:1 and 3:1 fraxetin-Fe complexes detected by Direct infusion ESI-QTOF MS analysis** (Figure 3). Theoretical isotope abundance was calculated using Data Analysis software from Bruker Daltonics

| Stoichiometry | Charge | | $R_h$ (nm) | Expected $\mu_{ep}$ (x10$^{-5}$ cm$^2$ V$^{-1}$ s$^{-1}$) | |
|---|---|---|---|---|---|
| **Fe:Fraxetin** | **$Fe^{3+}$** | **$Fe^{2+}$** | | **$Fe^{3+}$** | **$Fe^{2+}$** |
| 1:1 | +1 | 0 | 0.58[a] | +5.3 | 0 |
| 1:2 | -1 | -2 | 0.67[b1] / 0.72[b2] | -12.1[b1] / -10.7[b2] | -23.9[B1] / -21.9[b2] |
| 1:3 | -3 | -4 | 0.77[c] | **-27.2** | -32.9 |

**Table S2**. **Charge, hydrodynamic radius and electrophoretic mobility of Fe:Fraxetin complexes**. The charge of each complex was determined in the case of Fe(III) and Fe(II) and the hydrodynamic radius was determined on the simulated structure (Supplementary Fig. 5). Using the hydrodynamic radius and the charge, the theoretical electrophoretic mobility of each hypothetical complex could be determined using the OWO model and this latter value was compared to the experimental value of 27.02 × 10$^{-5}$ cm$^2$ s$^{-1}$ V$^{-1}$ obtained from the analysis of the Fe-Fraxetin mixture. Results show that only the 3:1 was in accordance with the experimental value and thus the other structures could be discarded.

| Metal | Molar masses | Isotope abundances (%) |
|---|---|---|
| Al | 26.9815 | 100 |
| Mn | 54.9380 | 100 |
| Fe | 55.9349 | 92 |
| | 53.9396 | 6 |
| | 56.9354 | 2 |
| Ni | 57.9353 | 68 |
| | 59.9308 | 26 |
| Cu | 62.9296 | 69 |
| | 64.9278 | 31 |
| Zn | 63.9291 | 48 |
| | 65.9260 | 28 |
| | 67.9248 | 19 |

**Table S3. Natural isotope abundances of the analyzed metals**

**Methods S1 Detailed protocols for pKa of Fraxetin determination and Effective charge of fraxetin and Fe(III)-fraxetin complex determination**

### p*Ka* of Fraxetin.

The determination of the acid constant of fraxetin was realised by capillary electrophoresis by analysing fraxetin at different pH values between 2.5 and 11. The calculated electrophoretic mobility, obtained from the transformation of the temporal electropherogram into an electrophoretic mobility distribution as described in (Chamieh *et al.*, 2015), was then plotted against the pH value and using the equation for a monoprotic acid (Poole *et al.*, 2004) (equation (1) below) a p$K_a$ value could be obtained as a fitting parameter. It is noteworthy that the second acidity of fraxetin was considered weak and not observed under the operating experimental conditions.

$$\mu_{ep} = \frac{\mu_{A^-} \times 10^{-pK_a}}{10^{-pK_a} + 10^{-pH}} \tag{1}$$

Were $\mu_{A-}$ is the electrophoretic mobility of fraxetin anion obtained at the plateau of the curve.

### Effective charge of fraxetin and Fe(III)-fraxetin complex.

The determination of the effective charge of fraxetin and Fe-fraxetin complexes was achieved by using the method based on the O'Brien-White-Ohshima model (OWO) (Ohshima, 2001; Ibrahim *et al.*, 2013):

$$\mu_{ep} = \frac{2\varepsilon_r \varepsilon_0 \zeta}{3\eta}\left[ f_1(\kappa R_h) - \left(\frac{ze\zeta}{k_B T}\right)^2 f_3(\kappa R_h) - \frac{m_- + m_+}{2}\left(\frac{ze\zeta}{k_B T}\right)^2 f_4(\kappa R_h) \right] \tag{2}$$

Where $\varepsilon_r$ is the relative electric permittivity of the media, $\varepsilon_0$ is the electric permittivity of vacuum ($8.85 \times 10^{-12}$ $C^2$ $J^{-1}$ $m^{-1}$), $\zeta$ is the zeta potential (V), $\eta$ is the viscosity of the media (Pa s), $\kappa$ is the Debye–Hückel parameter ($m^{-1}$) which represents the reciprocal thickness of the ionic cloud, $R_h$ is the hydrodynamic radius (m), $k_B$ is the Boltzmann constant ($1.38 \times 10^{-23}$ J $K^{-1}$), T is the temperature (K), $z$ is the charge number of the electrolyte ions, while $f_1$, $f_3$ and $f_4$ are functions of $R_h$ and are given by:

$$f_1(\kappa R_h) = 1 + \frac{1}{2\left[1 + 2.5/\{\kappa R_h\}\left(1 + 2e^{-\kappa R_h}\right)\right]^3} \tag{3}$$

$$f_3(\kappa R_h) = \frac{\kappa R_h\left(\kappa R_h + 1.3e^{-0.18\kappa R_h} + 2.5\right)}{2\left(\kappa R_h + 1.2e^{-7.4\kappa R_h} + 4.8\right)^3} \tag{4}$$

$$f_4(\kappa R_h) = \frac{9\kappa R_h\left(\kappa R_h + 5.2e^{-3.9\kappa R_h} + 5.6\right)}{8\left(\kappa R_h - 1.55e^{-0.32\kappa R_h} + 6.02\right)^3} \tag{5}$$

While $m_+$ and $m_-$ are dimensionless ionic drag coefficients, accessible from the limiting equivalent conductivities of the cation $\Lambda_+^0$ and the anion $\Lambda_-^0$ in the electrolyte and calculated using the following equation:

$$m_\pm = \frac{2\varepsilon_r\varepsilon_0 k_B T N_A}{3\eta z \Lambda_\pm} \tag{6}$$

The formalism suggested by Ohshima (Ohshima, 2001) requires the use of a symmetrical (z:z) electrolyte. If we consider the ammonium bicarbonate ($NH_4HCO_3$) buffer to be a 1:1 electrolyte with an ionic strength $I = C(NH_4^+)$, the dimensionless ionic drag coefficients of the cations and the anions can be calculated from the corresponding limiting ion conductivities which were found to be of $\Lambda_+^0$ = 68.16 S cm$^2$ mol$^{-1}$ for $NH_4^+$ at 25°C (calculated from data taken from (Tanaka & Hashitani, 1971) via the Einstein relationship) and $\Lambda_-^0$ = 44.71 S cm$^2$ mol$^{-1}$ for $HCO_3^-$ at 25°C (calculated from data taken from (Zeebe, 2011) via the Einstein relationship). From these limiting ion conductivities, the following ionic drag coefficients were obtained using equation (6): $m_+ = 0.189$ and $m_- = 0.288$.

This model can be applied for the effective charge determination of small ions, polyelectrolytes and nanoparticles (Ibrahim *et al.*, 2013) as long as the zeta potential $\zeta$ is lower than 100 mV and that $\kappa R_h$ is lower than 10 (Pyell *et al.*, 2009; Ibrahim *et al.*, 2012). Further, $\kappa$ is a function of the ionic strength of the media, $I$ (mol l$^{-1}$), and can be calculated using the following equation:

$$\kappa = \left( \sqrt{\frac{\varepsilon_0 \varepsilon_r k_B T}{2 N_A e^2 I 1000}} \right)^{-1} \tag{7}$$

Where $N_A$ is the Avogadro number (6.02 × 10$^{23}$ mol$^{-1}$), $e$ is the elementary electric charge (-1.6 × 10$^{-19}$ C) and 1000 is the conversion factor from l to m$^3$.

Next, analytical solutions of the OWO model (equation (2) and Supplemental Figure 12) relating the $\mu_{ep}$ to $\kappa R_h$ at different $\zeta$ values were plotted. Knowing, experimentally, the electrophoretic mobility $\mu_{ep}$ and the hydrodynamic radius of the analysed solute, $\zeta$ could be determined graphically and finally used for the calculation of the effective charge, $z_{eff}$, using the following equation (Makino & Ohshima, 2010):

$$z_{eff} = \frac{8\pi R_h^2 \varepsilon_r \varepsilon_0 \kappa k_B T}{ze^2} \sinh\left(\frac{ze\zeta}{2k_B T}\right) \times \left[1 + \frac{1}{\kappa R_h} \frac{2}{\cosh^2\left(ze\zeta/4k_B T\right)} + \frac{1}{\left(\kappa R_h\right)^2} \frac{8 \ln\left[\cosh\left(ze\zeta/4k_B T\right)\right]}{\sinh^2\left(ze\zeta/2k_B T\right)}\right]^{1/2}$$

(8)
